# Structure-function analysis of interallelic complementation in *ROOTY* transheterozygotes

**DOI:** 10.1101/2020.01.02.893016

**Authors:** Javier Brumos, Benjamin G. Bobay, Cierra A. Clark, Jose M. Alonso, Anna N. Stepanova

**Affiliations:** Department of Plant and Microbial Biology, Program in Genetics, North Carolina State University, Raleigh, NC 27695-7614, USA; Duke University NMR Center, Duke University Medical Center, Durham, NC 27710, USA; Department of Biochemistry, Duke University, Durham, NC 27710, USA; Department of Radiology, Duke University, Durham, NC 27710, USA

## Abstract

Auxin is a crucial plant growth regulator. Forward genetic screens for auxin-related mutants have led to the identification of key genes involved in auxin biosynthesis, transport, and signaling. Loss-of-function mutations in the genes involved in indole glucosinolate biosynthesis, a metabolically-related route that produces defense compounds from indolic precursors shared with the auxin pathway, result in auxin overproduction. We identified an allelic series of fertile, hypomorphic mutants for an essential indole glucosinolate route gene *ROOTY* (*RTY)* that show a range of high-auxin defects. Genetic characterization of these lines uncovered phenotypic suppression by *cyp79b2 b3*, *wei2*, and *wei7* mutants and revealed the phenomenon of interallelic complementation in several *RTY* transheterozygotes. Structural modeling of RTY shed light on the structure-to-function relations in the RTY homo- and heterodimers and unveiled the likely structural basis of interallelic complementation. This work underscores the importance of employing true null mutants in genetic complementation studies.

## Introduction

Forward genetic screens in the model plant Arabidopsis proved to be a powerful tool for identifying key players of the auxin biosynthesis, transport, and signaling machineries (Brumos et al., 2014; Kasahara, 2016; Zhao, 2014; Adamowski and Friml, 2015; Band et al., 2014; Estelle et al., 2011; Leyser 2018). In the past 30 years, the main components of these pathways have been uncovered and functionally characterized using a wide array of genetic, biochemical, and molecular approaches (Brumos et al., 2014; Kasahara, 2016; Zhao, 2014; Adamowski and Friml, 2015; Estelle et al., 2011; Leyser, 2018). In Arabidopsis, the predominant biologically active form of auxin, indole-3-acetic acid (IAA), is synthesized from the amino acid tryptophan via indole-3-pyruvic acid (IPyA) by the sequential action of two enzyme families, tryptophan aminotransferases, represented by TAA1, TAR1 and TAR2, and flavin-containing monooxygenases, represented by YUC1 through YUC11 (Supplemental Fig. S1A) (Stepanova et al., 2011; Mashiguchi et al., 2011; Won et al., 2011). Loss-of-function mutations that inactivate multiple *TAA1/TAR* or *YUC* genes lead to an array of auxin-deficient phenotypes, such as dwarfism, loss of apical dominance, reduced leaf venation, abnormal flower development and, in more severe cases, defective embryogenesis resulting in *monopteros*-like seedlings that lack hypocotyls and roots and develop a single cotyledon (Chen et al., 2014; Cheng et al., 2007; Stepanova et al., 2008; Stepanova et al., 2011). EMS alleles of *TAA1* that show mild auxin deficiency were originally isolated in genetic screens for reduced ethylene sensitivity (*wei8*) (Stepanova et al., 2008), loss of shade avoidance (*sav3*) (Tao et al., 2008), and resistance to auxin transport inhibitors (*tir2*) (Yamada et al., 2009), whereas higher order mutant combinations of the *TAA1* and/or *YUC* gene families were generated using T-DNA mutant alleles identified via reverse genetics (Mashiguchi et al., 2011; Stepanova et al., 2011). Furthermore, gain-of-function mutants of the *YUC1* gene have been found by screening an activation-tagged line collection and then recapitulated using transgenic approaches by expressing *YUC1* or other *YUC* genes under strong promoters (Zhao et al., 2001). *YUC* overexpression results in extreme overproduction of auxin and leads to plants with epinastic (i.e., downward curling) cotyledons and leaves, greater number of lateral and adventitious roots, and enhanced apical dominance in the inflorescences (Zhao et al., 2001).

Although the IPyA pathway is thought to be the main route of auxin synthesis in plants (Stepanova et al., 2011; Mashiguchi et al., 2011; Brumos et al., 2018), alternative branches may contribute to total IAA pools at least at some developmental stages, or in some tissues, or under specific environmental conditions. Loss-of-function mutations in several of the components of the parallel indole glucosinolate (IG) biosynthetic pathway, specifically *SUR2* (also known as *CYP83B1*, *RED1, RNT1* and *ATR4*)*, RTY* (*SUR1, ALF1, IVR*, and *HLS3*) and *UGT74B1* (*WEI9*), hyper-activate one of the putative IPyA-indepenent auxin biosynthetic branches by re-routing the pathway intermediate indole-3-acetaldoxime (IAOx, that is normally channeled into IG and camalexin production) into the biosynthesis of IAA (Supplemental Fig. S1A) (Barlier et al., 2000; Bak et al., 2001; Grubb et al., 2004; Mikkelsen et al., 2004; Nafisi et al., 2007; Müller et al., 2015; Mucha et al., 2019). IGs are secondary metabolites plants utilize to defend themselves against herbivorous pests. Knockouts of the IG biosynthetic genes *SUR2, RTY* and *UGT74B1* disrupt IG production and result in excess IAA accumulation. The high-auxin phenotypes of IG loss-of-function mutants are similar to that of *YUC* gain-of-function lines, with mutant plants displaying epinastic cotyledons and leaves and large numbers of adventitious and lateral roots (Bak et al., 2001; Grubb et al., 2004; Mikkelsen et al., 2004).

In this study, we describe an allelic series of hypomorphic mutants for one of the IG pathway genes, *RTY*. The RTY protein is a C-S lyase that catalyzes the conversion of S-(alkylacetohydroximoyl)-L-cysteine to thiohydroximate in the synthesis of IGs (Mikkelsen et al., 2004). Partial inactivation of this enzymatic step due to missense mutations in *RTY* leads to milder defects than previously reported for the seedling-lethal null mutants of this gene, with the developmentally stunted but fertile new alleles of *RTY* in our collection ranging in their severity from moderate to mild. Unexpectedly, genetic characterization of these lines uncovered several instances of interallelic complementation between missense mutants of *RTY*. Computational modeling of the RTY protein structure and mapping of RTY mutations onto the structural model of the RTY protein not only highlighted key residues essential for dimerization and binding of the substrate, but also shed light on structure-to-function relationships, revealing the likely structural basis of interallelic complementation. The combination of genetics and modeling employed in this work provided an effective, synergistic strategy to investigating the molecular basis of the complementation phenomenon in F1 transheterozygotes. Given the scarcity of reports describing interallelic complementation despite the ubiquitous use of allelic testing in the genetic characterization of mutants, this study serves as a cautionary tale that underscores the need of utilizing true null alleles of genes of interest in all complementation crosses.

## Results

### Characterization of a new allele of the RTY locus

Downward curling (i.e., epinasty) of cotyledons (Fig. 1A) and true leaves (Fig. 1B) is a characteristic feature of auxin biosynthesis mutants that produce excess IAA, such as gain-of-function mutants in the key IPyA pathway genes of the *YUC* family (Zhao et al., 2001) and loss-of-function alleles of the *SUR2* and *RTY* genes (Bak et al., 2001; Mikkelsen et al., 2004; Fig. 1A) disrupted in the IG branch of auxin biosynthesis (Supplemental Fig. S1A). Epinasty is, however, also observed in other classes of hormonal mutants, e.g., in the constitutive ethylene signaling mutant *ctr1* (Kieber et al., 1993; Fig. 1A) or brassinosteroid-insensitive mutant *bri1* (Clouse et al., 1996). In search for new ethylene and auxin mutants, we performed forward genetic screens that led to the isolation of dozens of lines with highly epinastic cotyledons, including multiple new alleles of known genes, such as *CTR1* and *SUR2*. One distinct subclass of mutants phenotypically resembled *rty* loss-of-function mutants (Mikkelsen et al., 2004) and *YUC* gene overexpression lines (Zhao et al., 2001), but showed varying degrees of epinasty and dwarfism in adults (see below). To gain insights into the nature of the affected gene(s), we performed phenotypic and genetic characterization of these lines.

**Figure 1.**
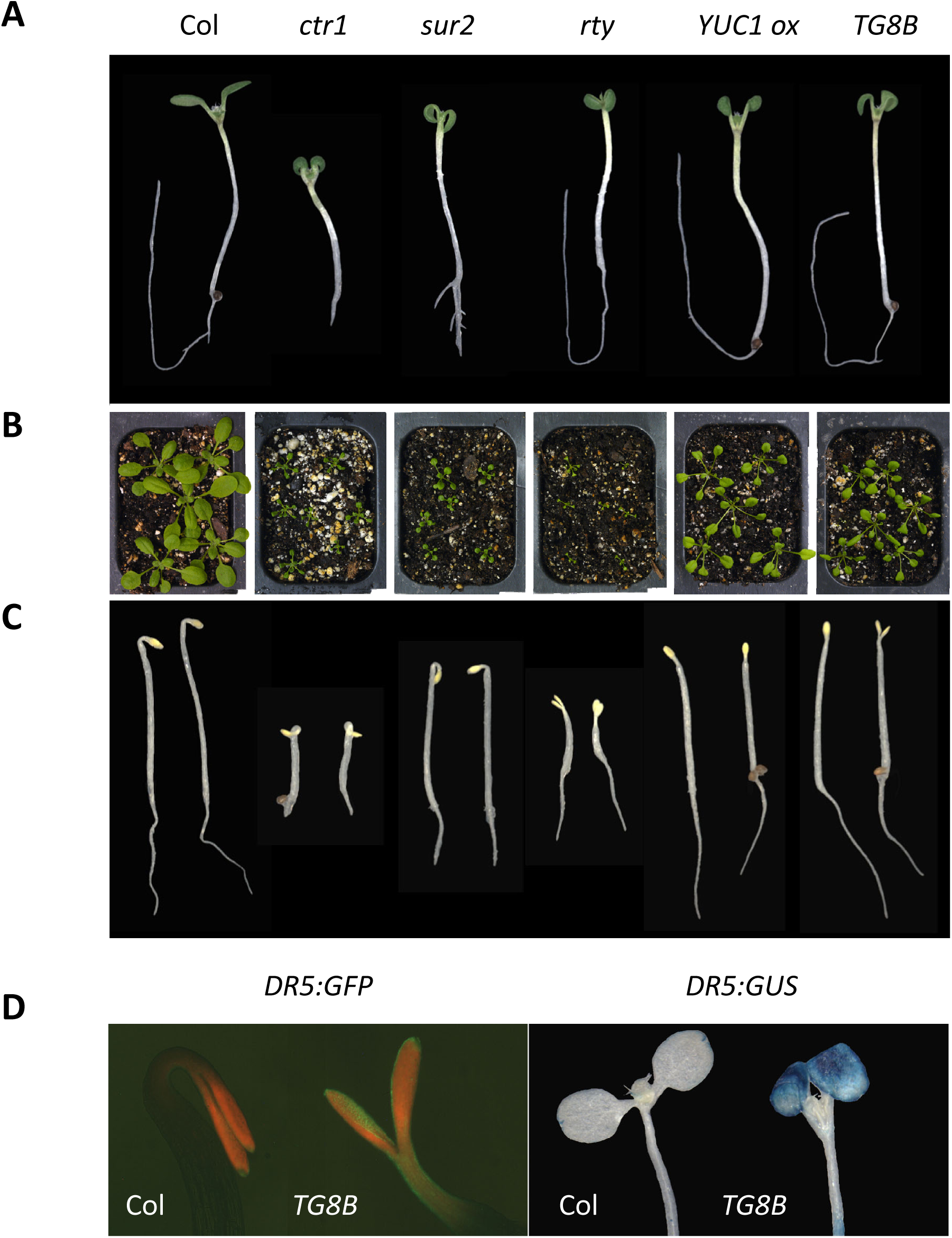
Phenotypes of mutants that share an epinastic cotyledon defect. Auxin overproducing and constitutive ethylene signaling mutants display profound cotyledon (A) and leaf (B) epinasty. (A) 8-day-old seedlings grown in plates for 3 days in the dark followed by 5 days in the light. (B) 4-week-old adults grown in soil. (C) 3-day-old plate-grown seedlings germinated in the dark. (D) Elevated *DR5* reporter activity in the new epinastic mutant *TG8B* relative to wild-type Col. 3-day-old dark-grown plate-germinated seedlings harboring *DR5:GFP* and 8-day-old plants grown in plates for 3 days in the dark followed by 5 days in the light harboring *DR5:GUS* are shown.

The first mutant of that subclass characterized was *TG8B* that showed characteristic cotyledon and leaf epinasty in light-exposed seedlings and soil-grown adults (Fig. 1A, B) and lacked apical hooks in etiolated seedlings (Fig. 1C), displaying phenotypes similar to that of *YUC1* overexpression lines (Zhao et al., 2001) and *rty* loss-of-function mutants (Mikkelsen et al., 2004). The expression of the auxin-response reporters *DR5:GFP* and *DR5:GUS* in this line was dramatically enhanced, consistent with the increased auxin activity in *TG8B* (Fig. 1D) comparable to that of other auxin overproducers (Supplemental Fig. S1B, S1C). Genetic analysis of *TG8B* in the F1 generation of crosses with Col *DR:GFP* and Col *DR5:GUS* (reporter introgression backcrosses) and L*er* (mapping) revealed recessive nature of the mutation (Table 1), suggesting that the *TG8B* phenotype is unlikely to be caused by a direct activation of one of the *YUC* genes. Accordingly, we next tested *TG8B* for potential allelism with known recessive auxin overproducing mutants, *sur2* and *rty*. The F1 progeny of the cross between *TG8B* and *sur2 DR5:GUS*, a T-DNA null mutant with the introgressed auxin reporter (Ulmasov et al., 1997; Stepanova et al., 2005), was phenotypically wild-type (Table 1), suggesting that the two mutants are not allelic. In the complementation test with *rty*, we utilized the fertile *rty wei2-1* (Stepanova et al., 2005) double mutant combination as a parent, because the *rty* single mutant is lethal. As in the case with *sur2*, the F1 generation of reciprocal crosses between *TG8B* and *rty wei2-1* was phenotypically wild-type (Table 1). This suggested that *TG8B* may not be allelic to *rty*. We therefore performed PCR-based mapping on the phenotype-selected F2 population derived from a cross between *TG8B* (Col) and L*er*. To our surprise, despite the complementation test results, *TG8B* was found to be tightly linked to the *RTY* locus (*At2g20610*), with 0 recombinants observed in 58 chromosomes for the *F26H11-1* indel PCR marker located at 9.065 Mb on chromosome 2 next to the *RTY* gene (8.878 – 8.880 Mb) (Supplemental Table S1).

**Table 1.** Crosses of new epinastic mutants.

| Cross (female x male) | Phenotype in F1 |
| --- | --- |
| ***TG8B*** selfed<br>***Col DR5:GFP x TG8B***<br>***Col DR5:GUS x TG8B***<br>***Ler x TG8B***<br>***sur2 DR5:GUS x TG8B***<br>***wei2-1 rty x TG8B***<br>***TG8B DR5:GFP x wei2-1 rty*** | Weak *rty*-like (see Fig. 3, 4)<br>WT-like<br>WT-like<br>WT-like<br>WT-like<br>WT-like<br>WT-like (see Fig. 4) |
| ***TH5H*** selfed<br>***Col x TH5H***<br>***sur2 DR5:GUS x TH5H***<br>***Ler x TH5H***<br>***wei2-1 rty x TH5H*** | Moderate *rty*-like (see Fig. 3, 4)<br>WT-like<br>WT-like<br>WT-like<br>WT-like (see Fig. 4) |
| ***TF5D*** selfed<br>***Col DR5:GFP x TF5D***<br>***Ler x TF5D***<br>***wei2-1 rty x TF5D DR5:GFP*** | Moderate *rty*-like (see Fig. 3, 4)<br>WT-like<br>WT-like<br>Weak *rty*-like (see Fig. 4) |
| ***TD11B*** selfed<br>***Col DR5:GFP x TD11B***<br>***Ler x TD11B***<br>***TD11B x sur2 DR5:GUS***<br>***TD11B DR5:GFP x wei2-1 rty*** | Very weak *rty*-like (see Fig. 3, 4)<br>WT-like<br>WT-like<br>WT-like<br>Very weak *rty*-like (see Fig. 4) |
| ***TJ12H*** selfed<br>***Col DR5:GFP x TJ12H***<br>***sur2 DR5:GUS x TJ12H*** | Very weak *rty*-like<br>WT-like<br>WT-like |
| ***IK6G12*** selfed<br>***Col x IK6G12***<br>***wei2-1 rty x IK6G12*** | Moderate *rty*-like (see Fig. 3, 4)<br>WT-like<br>WT-like (see Fig. 4) |
| ***CS878294*** selfed<br>***CS878294 x wei2-1 rty*** | Strong *rty*-like (see Fig. 3, 4)<br>Strong *rty*-like (see Fig. 3, 4) |

To determine whether *TG8B* is, indeed, an allele of *RTY*, we developed a set of five primer pairs that amplify five overlapping gene fragments and cover the entire *RTY* gene (Fig. 2A, Supplemental Table S1). We PCR-amplified *RTY* from *TG8B*, Col, and *rty* genomic DNA and performed a CEL1 assay on each mutant against the Col *RTY* sample. CEL1 is a crude mixture of DNases from celery that preferentially cut mismatched bases in a DNA heteroduplex formed by annealing of denatured PCR-amplified wild-type and mutant samples (Till et al., 2004). CEL1 assay thus allows for the identification of polymorphisms in an amplified mutant gene fragment with respect to the corresponding wild type fragment, when the products of the CEL1 digestion reaction are separated by regular agarose gel electrophoresis (Till et al., 2004). Polymorphisms were detected by CEL1 assay for both mutants: in fragment 3 for *rty* and fragment 4 for *TG8B* (Fig. 2A). Presence of mutations was then confirmed by Sanger sequencing. *rty* was found to harbor a C893T transition in the 2^nd^ exon of *RTY* genomic DNA that results in a P213S amino acid substitution, whereas *TG8B* had a polymorphism later in the gene, in the 4^th^ exon, mutating G1412A in the genomic DNA and thus replacing an aspartate by an asparagine (D315N) in the RTY protein. The two mutations affect strongly- and moderately-conserved amino acids (Fig. 2B) and [based on the strength of mutant phenotypes] result in the complete or partial loss of *RTY* function in *rty* and *TG8B*, respectively. To further confirm the molecular identity of *TG8B* as an allele of *RTY*, we transformed the mutant with a cDNA construct of *RTY* driven by the *35S* promoter and observed nice complementation of the cotyledon and leaf epinasty and dwarfism rescue (Fig. 2C).

**Figure 2.**
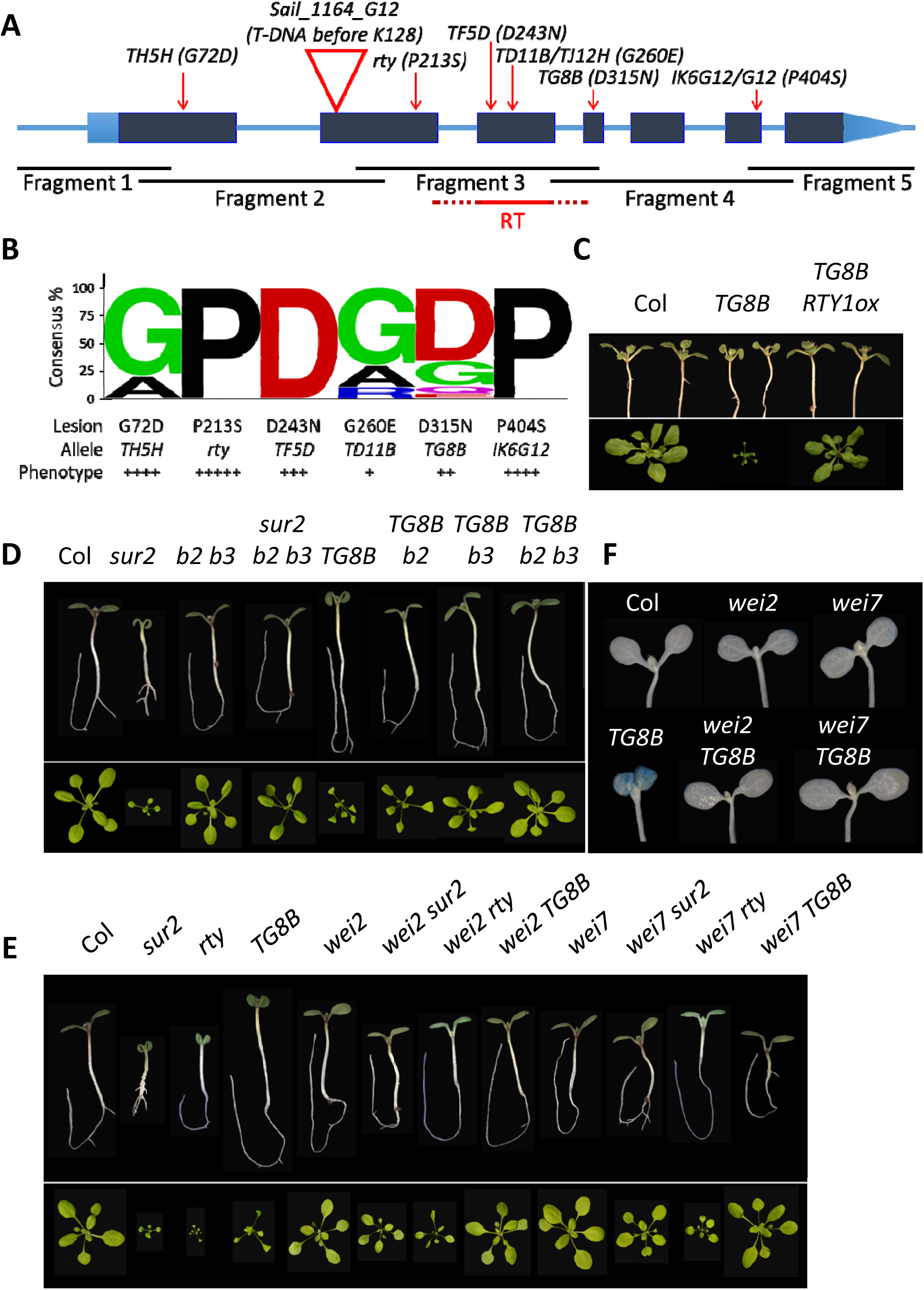
Identification of *TG8B* as a hypomorphic allele of *RTY*. (A) A schematic of the *RTY* gene displaying its exon (boxes) and intron (inner lines) structure. Dark blue boxes correspond to coding regions, whereas light blue boxes represent the 5’ and 3’ UTRs. Locations of mutations are marked with red arrows (substitutions) and triangles (T-DNA insertions) and the nature of mutations is indicated next to the mutant line name. Overlapping PCR fragments amplified for CEL1 assay and sequencing are shown as black lines below the gene model. The fragment shown in red corresponds to the reverse transcription product used in the quantification of *RTY* gene expression in Figure S5. (B) Multiple Sequence Alignment among RTY-like proteins was performed to analyze the conservation of the mutated amino acids of RTY. The sequence logo reveals a correlation between amino acid conservation and the severity of missense mutant phenotypes, with the weaker alleles of *RTY*, *TD11B* and *TG8B*, showing the least amino acid conservation, and the strongest allele, *rty*, showing full conservation. (C) Complementation of *TG8B* with a cDNA construct of *RTY*. 10-day-old T2-generation seedlings grown in plates for 3 days in the dark followed by 7 days in the light and 4-week-old soil-grown adults are shown next to age-matched control plants. (D, E, F) *cyp79b2 b3* double mutant (D), *wei2* and *wei7* (E, F) can suppress the high-auxin phenotypes of *TG8B.* 8-day-old seedlings grown in plates for 3 days in the dark followed by 5 days in the light and soil-grown 4-week-old adults are shown. (F) GUS staining in the *TG8B wei2* and *TG8B wei7* double mutants is reduced relative to *TG8B* itself.

Consistent with *TG8B* being an allele of *rty*, double mutant *cyp79b2 cyp79b3* that is disrupted in the conversion of tryptophan to IAOx (Supplemental Fig. S1A) and thus devoid of the indole-glucosinolate pathway precursor IAOx (Zhao et al., 2002), could fully suppress the high-auxin phenotypes of *TG8B* (Fig. 2D). Single *cyp79b2* or *cyp79b3* mutants could only partially rescue *TG8B* (with the *cyp79b3* being the more effective suppressor of the two), a finding that is in agreement with the partial functional redundancy of *CYP79B2* and *CYP79B3* (Zhao et al., 2002). The phenotypic complementation of *TG8B* by mutations in the first committed step of the IAOx route catalyzed by the CYP79B family is in agreement with a previous report that showed the ability of *cyp79b2 cyp79b3* to also revert *sur2* (Stepanova et al., 2011; Fig. 2D), another auxin overproducing mutant that maps to the IG branch of the auxin pathway downstream of the *CYP79B* step but upstream of *RTY* (Supplemental Fig. S1A) (Bender and Celenza, 2009), back to wild-type. Furthermore, *wei2* and *wei7*, the anthranilate synthase alpha and beta subunit mutants defective in the tryptophan biosynthesis pathway (Supplemental Fig. S1A) that were previously shown to suppress the high-auxin phenotypes of *sur2* and *rty* by limiting the availability of the auxin precursor tryptophan (Stepanova et al., 2005), could also suppress *TG8B*, resulting in improved plant growth (Fig. 2E) and normalized auxin response (Fig. 2F).

### Possible genetic mechanisms of RTY complementation in F1

We next decided to investigate why the F1 generation of the cross between the classical *rty* mutant (King et al., 1995) and the new allele, *TG8B*, showed full complementation. We could envision two possibilities. One possible scenario explaining the wild-type morphology of the *rty/TG8B* transheterozygote is that the *rty* phenotype might have been suppressed by one copy of the *wei2-1* mutation, since a *rty wei2* double mutant was utilized for this cross and, therefore, the F1 was genotypically *rty/TG8B wei2/+*. This is, however, an unlikely possibility, as two copies of the *wei2-1* mutant allele are required for the suppression of the high-auxin defects of *sur2* and *rty* (Stepanova et al., 2005) or of *TG8B* (Fig. 2E, 2F), and the sesquimutants *TG8B/TG8B wei2/+, rty/rty wei2/+* (Supplemental Fig. S2A, S2B) and *sur2/sur2 wei2/+* (Supplemental Fig. S2C) are phenotypically similar to *TG8B*, *rty*, and *sur2*, respectively. The second possible scenario is that the RTY protein may function as a dimer (or a higher order multimeric complex) and thus in the F1 transheterozygote (that harbors an interallelic combination of two different *RTY* alleles, *rty* and *TG8B*) forms a fully functional heteromer. In other words, if the two RTY monomers in the F1 are defective in different domains of the protein, they may be able to complement one another by supplying the missing activity to the dimer/oligomer. In fact, RTY belongs to a large family of C-S lyases/transaminases/alliinases several of which have been shown to function as dimers or tetramers (Nock and Mazelis, 1987; John, 1995; Breitinger et al., 2001). The closest related Arabidopsis protein in the database for which the crystal structure is available, At2g22250, an aspartate/prephenate aminotransferase MEE17, is known to form dimers (Holland et al., 2018). Consistent with the possibility of transheterozygote complementation, the amino acid substitutions in *rty* and *TG8B* map to different domains of the protein (see below), suggesting that they may compromise different activities. Thus, transheterozygote complementation in *RTY* is a plausible explanation.

### Interallelic complementation in the RTY locus is not uncommon

To further explore the basis of mutant phenotype complementation in the *RTY* transheterozygote, we analyzed several additional putative *rty* alleles identified in our mutant screen. *TD11B, TF5D*, and *TH5H* show varying degree of cotyledon and leaf epinasty in light-grown plants (Fig. 3A, B) and lack apical hooks in dark-grown seedlings (Fig. 3C), as do *rty* and *TG8B*. Adult phenotypes range in severity in terms of rosette size and fertility, with *TD11B* being the weakest of all mutants, and *TH5H* being the strongest in this subset, but not as severe as *rty* itself (Fig. 3B, D). It is not surprising that neither of the novel mutants is as strong as the classical *rty* allele (King et al., 1995), as only fertile mutants have been recovered from our epinastic cotyledon genetic screens. In fact, several additional *rty*-like lines were initially selected in seedlings but lost during propagation due to their lethality in young adults.

**Figure 3.**
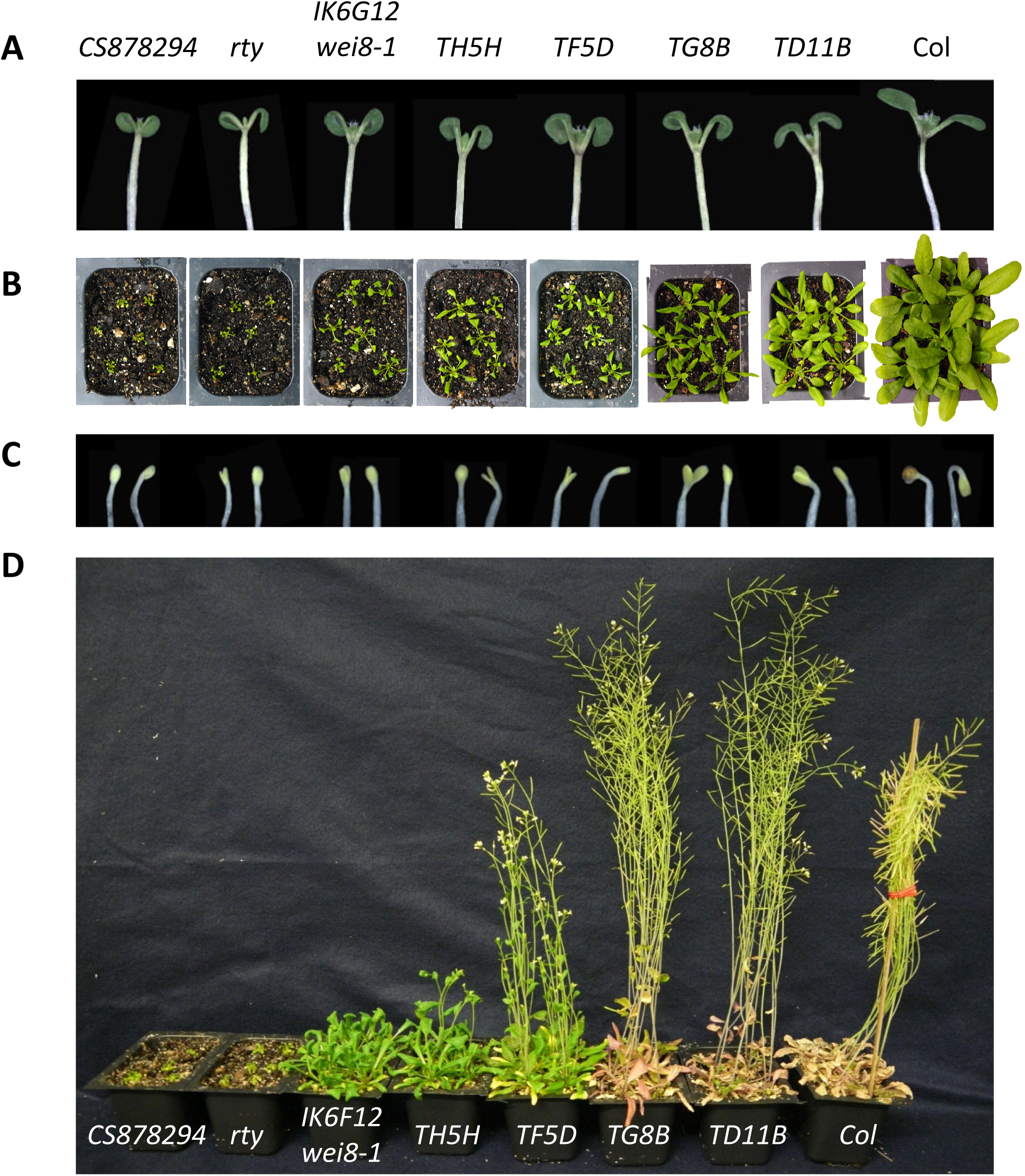
An allelic series of hypomorphic to null alleles of *RTY*. (A) 8-day-old seedlings grown in plates for 3 days in the dark followed by 5 days in the light. (B) 5-week-old adults grown in soil. (C) 3-day-old plate-grown seedlings germinated in the dark. (D) 12-week-old adult plants grown in soil. Note that the control wild-type Col plant was staked and its inflorescences tied and folded to prevent seed loss during harvesting.

Backcrosses of *TD11B, TF5D*, and *TH5H* to wild-type Col and L*er* revealed that all three mutants are recessive (Table 1). CEL1 assay, sequencing and/or mapping of these mutants were performed and demonstrated that all of these lines are, indeed, new alleles of *rty* (Fig. 2A). *TD11B* harbors a G1161A mutation in the 3^rd^ exon of *RTY* genomic DNA that alters glycine 260 to glutamic acid in the RTY protein. Rough mapping of *TD11B* (Col) in the F2 generation of a cross to L*er* supports the notion that *TD11B* is an allele of *RTY*, as 0 out of 40 chromosomes were recombinant with the *F26H11-1* indel PCR marker (Supplemental Table S1) that is less than 200Kb away from the *RTY* locus. *TF5D* shows a CEL1-assay detectable polymorphism in the fragment 3 of *RTY* (Fig. 2A). Sequencing of this fragment in the *TF5D* mutant background has uncovered a G1109A nucleotide substitution in *RTY* genomic DNA that corresponds to a D243N mutation in the 3^rd^ exon. Furthermore, another recessive *rty*-like mutant, *TJ12H*, that harbors an identical mutation to that of *TD11B* (Fig. 2A) but has been isolated from a different EMS family and thus has arisen independently from *TD11B*, is phenotypically indistinguishable from *TD11B* (Table 1), further suggesting that the high-auxin phenotypes of both of these mutants are a consequence of the D243N amino acid substitution in the RTY protein and not some other mutation. Finally, the *TH5H* mutant has been found to harbor a G215A missense mutation in the 1^st^ exon of *RTY* genomic DNA, resulting in a G72D amino acid substitution (Fig. 2A). Taken together, these results demonstrate that an allelic series of the *RTY* locus has been identified. Not surprisingly, the severity of the high-auxin phenotypes in *RTY* mutants (Fig. 3) correlates with the degree of conservation of the specific amino acid affected by the mutation (Fig. 2B).

To test the ability of the new *rty* alleles to form functional heteromers in F1 transheterozygotes, we intercrossed all of the available *RTY* locus mutants, i.e. *rty* (in the *wei2-1* background for fertility reasons)*, TG8B, TD11B, TF5D*, and *TH5H*, and examined the morphology of the F1 generation (Fig. 4). Remarkably, some but not all interallelelic combinations were able to complement each other, potentially offering insights into the structure-function relations in RTY heteromers. For example, the *TG8B* amino acid substitution mutant was able to complement all other missense alleles of *RTY* to wild-type level (Fig. 4), demonstrating full interallelic complementation in F1 transheterozygotes. Good complementation was also observed between two stronger *RTY* alleles, the classical *rty* and *TH5H*, whereas no or only very weak complementation was seen between *rty* or *TH5H* and the moderate allele *TF5D* (Fig. 4). Several *RTY* allele intercrosses (e.g., *rty/TD11B*, *TH5H/TD11B*, and *TF5D/TD11B* F1 transheterozygotes) displayed phenotypes less severe than that seen in the parents (in this case, *TD11B*) (Fig. 4, Supplemental Fig. S3), yet the morphology of these F1s was also distinct from that of wild-type plants, suggesting partial complementation.

**Figure 4.**
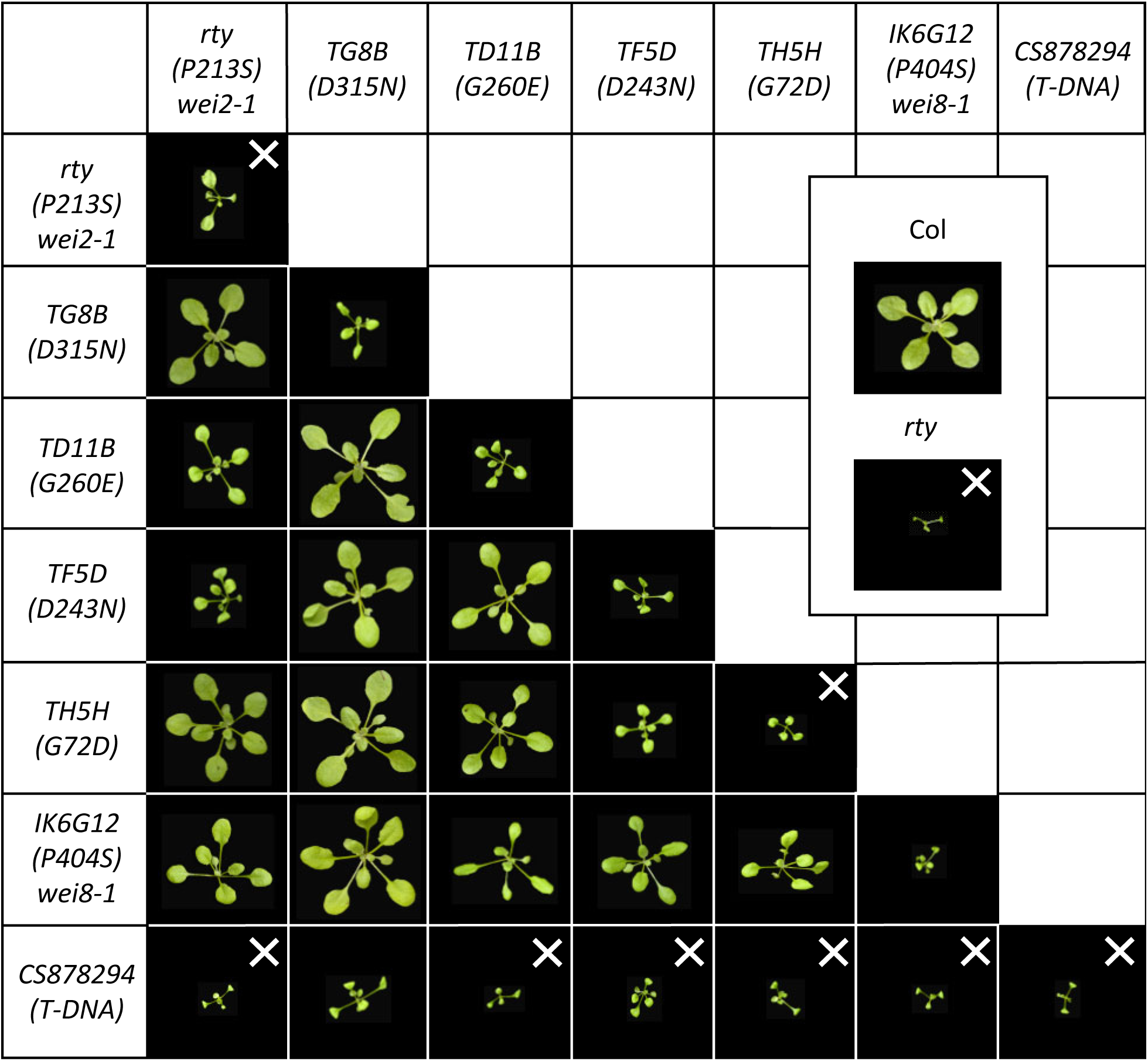
Interallelic combinations of *RTY* mutants. Phenotypes of 4-week-old soil-grown F1 and parental plants are shown. “X” marks the lines that failed to set seeds in two independent F1 propagation experiments. Note that the *rty wei2-1* and *TH5H* parental lines are able to set a limited amount of seeds in some trials.

In summary, we found that the phenomenon of interallelic complementation is not uncommon in the *RTY* gene, with all missense mutations examined complementable by at least one other missense allele. More importantly, although we found that the strength of the homozygous mutants correlated well with the degree of conservation of the affected amino acid, there was no clear correlation between the strength of the individual alleles in the homozygous plants and the severity of the defects in the heteroallelic combinations in the F1s of the intercrosses. This suggests that in some heteroallelic combinations specific structural changes induced by the monomer of one allele can compensate for the changes imposed by a different allele in the second monomer. To examine this possibility, we determined the predicted structural changes of several homo- and heteroallelic combinations of RTY.

### Structural details inferred from molecular models of RTY

We next employed molecular modeling to provide structural reasoning for the ability of transheterozygous mutations in *RTY* to restore the RTY activity and thus lead to a phenotype that is milder than either of the homozygous parents. A three-dimensional (3D) structural model of dimeric RTY was generated using the crystal structure of the closest RTY homolog from *Arabidopsis thaliana*, the aforementioned aminotransferase MEE17 (PDB code: 5WMH) (Holland et al., 2018) (Supplemental Fig. S4). The mutations described above were then mapped onto the wild-type dimeric RTY model (Fig. 5, Table 2), landing in varied locations throughout the dimeric structures. In particular, the D315 residue (mutated in *TG8B*) is located directly in the dimerization interface, suggesting that an amino acid substitution at this site may affect the efficiency of RTY protein dimerization. This site also falls within 8Å from the PLP cofactor and substrate binding pocket of the other monomer (across the dimerization interface) and lines the binding pocket entrance. To experimentally test the possible involvement of the D315N in the dimer formation, wild-type RTY (WT) and the D315N (TG8B) mutant were expressed in a heterologous system as part of bait and prey hybrid proteins in yeast, one fused with a transcriptional activation domain (AD) and another with a DNA binding domain of GAL4 (BD) (Supplemental Fig. S5). We observed that the activity of the LacZ reporter in plates was dramatically reduced when both the bait and the prey carry the D315N mutation (TG8B + TG8B), whereas a nearly wild-type level of the reporter activity was seen for the interallelic combination with WT RTY (WT + TG8B) (Supplemental Fig. S5A). Quantification of the beta-galactosidase activity in yeast using liquid assays revealed that the interallelic WT-TG8B combination yielded about 80% of the WT-WT activity (Supplemental Fig. S5B), consistent with the recessive nature of *TG8B* in plants. These results provide a plausible explanation for the observed interallelic complementation by D315N in pairwise combinations with other missense mutations of *RTY* in *Arabidopsis*. Specifically, the phenotype of the homozygous D315N mutant is the result of its inability to form homodimers, but dimer formation can be largely restored if only one of the monomers has the D315N mutation. The fact that all interallelic combinations of *TG8B* (D315N) with other missense alleles in our collection display wild-type morphology suggests that those other alleles of *RTY* do not interfere with the dimerization process. These results also imply that the D315N mutation does not directly affect the catalytic activity of the protein besides preventing the formation of functionally active RTY homodimers. Therefore, if a heterodimer forms between D315N and a catalytically impaired monomer such as D243N (see next), the resulting dimer should still be able to carry out the C-S lyase reaction (due to the catalytic functionality of D315N).

**Figure 5.**
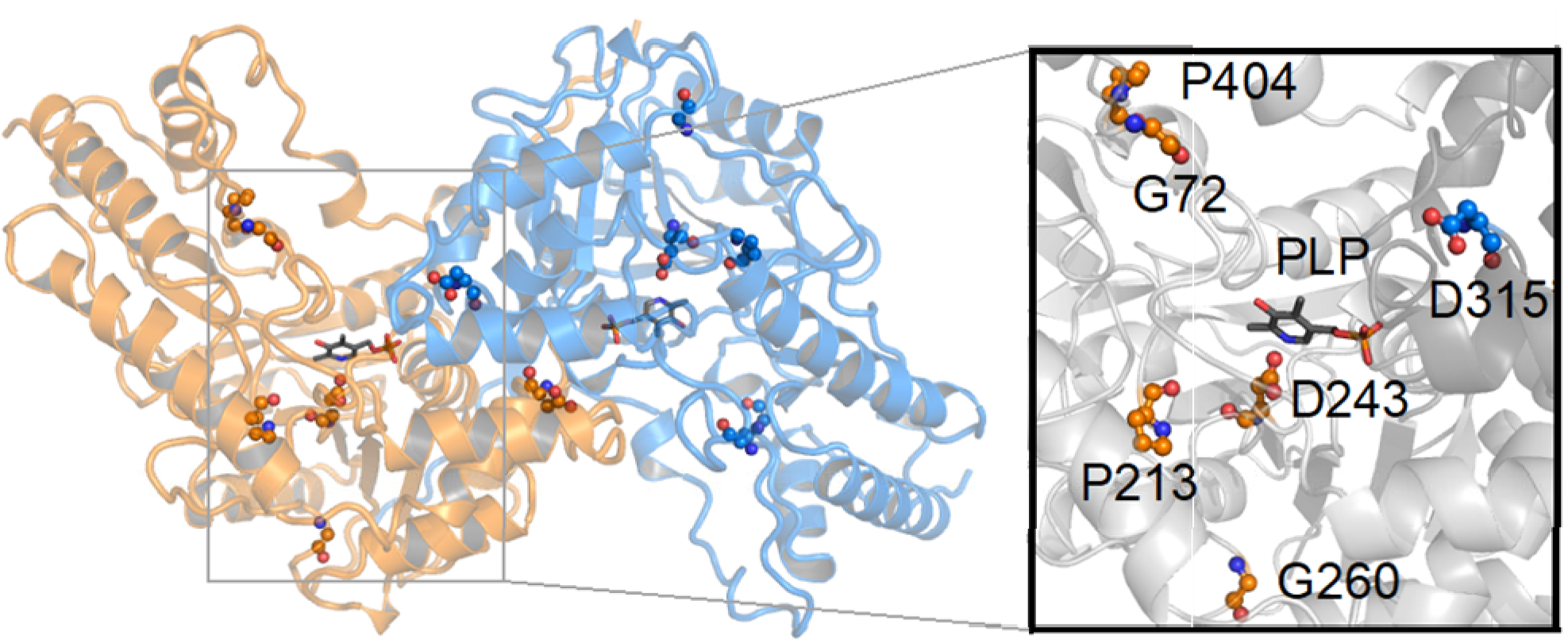
Dimeric model of RTY. Location of mutated residues and detailed view of the PLP binding pocket in the RTY protein. The mutations are labeled and shown in sphere-and-stick format on a background of the protein structure shown in cartoon format. One monomer is colored orange while the other is colored blue. PLP ligand, shown in stick-and-ball format, is included for clarity. The inset provides a detailed view of the substrate binding pocket and the location of the mutations that line the entrance and bottom of the PLP binding pocket. The surface is colored grey, while the mutations are colored orange and blue and shown in sphere-and-stick format.

**Table 2.**
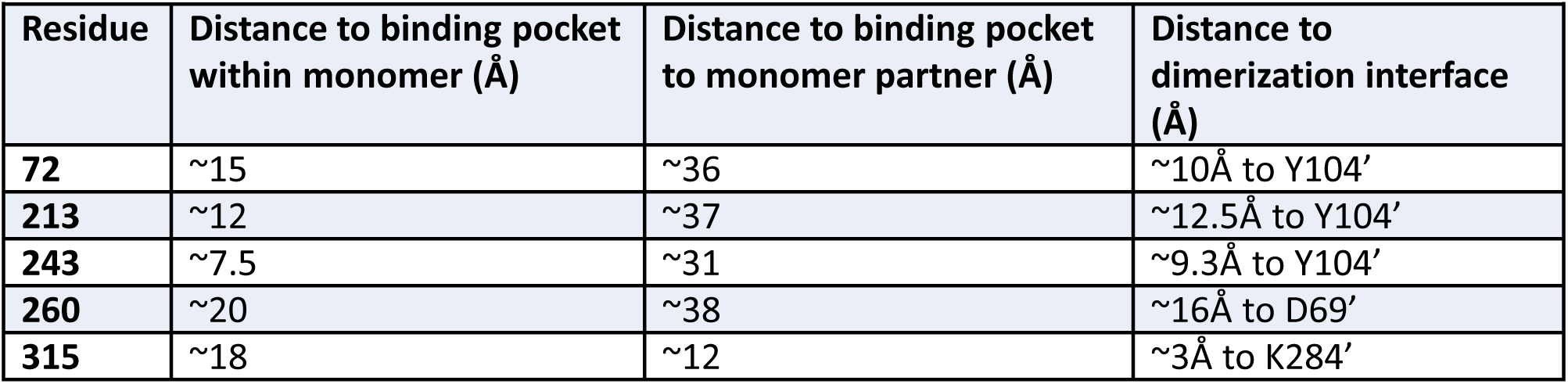
RTY Mutations and the distances to the binding pocket and dimerization interface.

| Residue | Distance to binding pocket within monomer (Å) | Distance to binding pocket to monomer partner (Å) | Distance to dimerization interface (Å) |
| --- | --- | --- | --- |
| **72** | ~15 | ~36 | ~10Å to Y104' |
| **213** | ~12 | ~37 | ~12.5Å to Y104' |
| **243** | ~7.5 | ~31 | ~9.3Å to Y104' |
| **260** | ~20 | ~38 | ~16Å to D69' |
| **315** | ~18 | ~12 | ~3Å to K284' |

The second *RTY* mutation for which the structural model provides a likely explanation is D243N (defective in *TF5D*), for which the side-chain of D243 falls in the binding pocket of the pyridoxal phosphate (PLP) cofactor (between 1-2 Å) (Fig. 5) and thus most likely affects the ability of RTY subunits to recruit PLP and the substrate, potentially resulting in reduced activity of mutant RTY. Consistent with this explanation, the heteroallelic combination D315N/D243N shows a wild type plant phenotype, as we anticipate that this allelic combination would be able to form dimers allowing for the catalytic domain of D315N to function even if the catalytic domain of D243N is largely inactive.

Other RTY mutations, G72D (*TH5H*), P213S (*rty*), and G260E (*TD11B/TJ12H*), map away from the catalytic pocket or the dimerization interface and therefore are harder to interpret with respect to the mutant defects they trigger (Fig. 5, Table 2). In these cases, we wondered whether the phenotypes of different interallelic combinations could provide some clues on the structure-function relationship of these mutations. The wild-type phenotype of the interallelic combination between G72D and the dimerization mutant D315N suggests that the G72D is unlikely to directly affect dimerization. On the other hand, the strong transheterozygous mutant defects of the G72D and the PLP-binding-impaired D243N combination would suggest that these two mutants may affect the same function of the protein, in other words, the PLP binding and/or substrate recruitment. In fact, this is not so far-fetched based on the structure analysis, as the residue G72 is solvent exposed and lines the entrance to the PLP binding channel in our structural model (Fig. 5). The positions of the two other missense mutations, P213S (that is found in our strongest allele, *rty*) and G260E (seen in the weakest allele in the series, TD11B), in the 3D model do not provide strong clues on the possible structure-function alterations in these mutants, as they are located distal to the PLP binding pocket. P213 and G260 occupy a center of mass (COM - a point representing the mean position of the matter in a protein), space ∼7 and 13Å away from D243 (the residue in the substrate binding pocket likely implicated in PLP binding), respectively (Fig. 5).

### Experimental testing of the 3D structural model of RTY

To test the potential capacity of the computational model of RTY to explain the phenotypes (complemented or not complemented) of F1 transheterozygotes, we have mapped the amino acid substitution of another novel *RTY* allele, *IK6G12*, which has been recently identified in our laboratory from another genetic screen in the *wei8-1* mutant background. From prior studies we knew that *wei8-1* is deficient in the tryptophan aminotransferase activity in the parallel IPyA branch of auxin biosynthesis (Supplemental Fig. S1) and this mutation does not suppress the epinastic cotyledon and leaf phenotypes of *rty* and *sur2* (Stepanova et al., 2008). *IK6G12* (*wei8-1*) is a moderately strong recessive allele of *RTY* (Fig. 3) that harbors a C1890T missense mutation in the sixth exon that alters proline 404 to serine (Fig. 2A, B). Based on our 3D model, P404 is located ∼19 Å from the PLP binding pocket, ∼40 Å from the PLP binding pocket within the other monomer, and ∼16 Å from the nearest residue in the dimerization interface (Y104) (Fig. 5, Table 2). Thus, the structural effects of P404S are likely to be indirect through creating conformational changes to the entrance of the PLP binding pocket or alteration to the dimerization interface and, therefore, should be functionally different than the structural consequences of all other missense mutants characterized in our study. Accordingly, we predicted that the *IK6G12* allele should be able to, at least in part, complement all of the available missense mutants. In fact, when we performed crosses between *IK6G12* and other alleles of *RTY* to experimentally test our predictions, we found that indeed *IK6G12* was able to fully complement *TG8B* and partially all other missense mutants in the F1 transheterozygotes (Fig. 4). These observations support the idea that interallelic complementation is possible for mutant combinations that disable functionally non-overlapping protein domains. Our findings are also consistent with the notion that lack of complementation may be indicative of the two allelic mutations affecting domains of similar functions.

As the genetic analyses and computational modeling of RTY described above strongly indicate that the use of missense mutations in complementation tests can be misleading, we decided to also utilize a true null allele of *RTY*. Our expectation was that a mutant that makes no full-length RTY protein should be unable to complement any of the amino acid substitution alleles of *rty*. We identified a previously uncharacterized insertional allele of *RTY*, *Sail_1164_G12* (CS878294), that harbors a T-DNA at the beginning of the second exon of *RTY* immediately upstream of the nucleotide 638 in the *RTY* genomic DNA (Fig. 2A). This T-DNA line can still make partial 3’ *RTY* mRNA, as detected by qRT-PCR with primers that anneal downstream of the insertion site (Supplemental Fig. S6), but the levels of the 3’ product are reduced more than three fold. The corresponding truncated protein, if produced, should be non-functional as it is expected to lack at least the first 207 amino acids, since the first available methionine downstream of the T-DNA insertion site in frame with the RTY protein is M208. Based on our mutant analysis, even a single amino acid substitution in the N-terminus, G72D (TH5H), leads to severe functional consequences, and the deletion of the entire N-terminal part of the protein would remove significant portions of the dimerization interface (86%) as well as nearly half of the PLP binding pocket. It is highly unlikely that this protein can fold correctly, let alone be able to dimerize and function.

Consistent with the notion that the *Sail_1164_G12* allele is null, the mutant was unable to rescue any of the missense *RTY* alleles in interallelic combinations (Fig. 4). This lack of complementation is not simply due to a more severe phenotype of *Sail_1164_G12* compared to the other alleles of *RTY* employed in this study, as the strength of this T-DNA mutant is similar to that of the classical missense allele, *rty*, utilized herein (Fig. 3). Since *Sail_1164_G12* is an exonic insertional mutant that cannot produce full-length *RTY* RNA, in F1 transheterozygotes this mutant should not be able to form RTY protein heteromers and should only produce homomeric mutant RTY protein complexes. Consistent with that expectation, for the majority of *RTY* mutants, the severity of the phenotypes in the F1 generation of inter-allelic crosses with the T-DNA allele of *RTY* was found to be enhanced with respect to that of the homozygous missense mutant parents (Fig. 4), suggesting that these interallelic *RTY* mutants may be haploinsufficient. Surprisingly, the *TG8B/Sail_1164_G12* F1 transheterozygotes were phenotypically indistinguishable from *TG8B* itself (Fig. 4), suggesting that in this mutant the number of the oligomers formed may not be rate-limiting despite the reduced ability of this mutant to dimerize relative to wild-type RTY.

## Discussion

### Genetic analysis

In this study, we identified and characterized an allelic series of missense mutants of *RTY* derived from EMS screens for epinastic ethylene- and auxin-related mutants. While complementation analyses combined with phenotyping proved to be a very useful and efficient technique for sub-grouping *CTR1* and *SUR2* mutant alleles identified in these screens, complementation tests largely failed when trying to categorize new alleles of *RTY* using a previously characterized presumed null allele, *rty* (King et al., 1995). In contrast, standard molecular biology approaches, including classical PCR-based mapping, CEL1 assay, and Sanger sequencing, were successful at identifying *RTY* as the causal gene for the six novel amino acid substitution mutants. Mutant complementation by a cDNA construct confirmed the identity of *RTY* as the gene responsible for *TG8B* epinasty. Genetic suppression of the high-auxin phenotypes of *TG8B* by the double *cyp79b2 cyp79b3* mutant provided additional support for attributing the auxin phenotype of this novel mutant to a defect in the IG branch of auxin biosynthesis. This result is consistent with the previously described ability of *cyp79b2 cyp79b3* to suppress *sur2* (Stepanova et al., 2011), i.e. another IG mutant, and further confirms the role of *RTY* in the IAOx-mediated auxin production. Likewise, the elevated levels of *DR5* reporter activity and the suppression of *TG8B* phenotypic defects by *wei2* and *wei7* mutants (Stepanova et al., 2005) deficient in the rate-limiting biosynthetic step of the auxin precursor tryptophan are also in agreement with the notion that the phenotypic defects of this *RTY* mutant arise from auxin overproduction.

In contrast with the problems encountered when employing missense *RTY* mutants in complementation testing, utilization of the true null T-DNA allele of *RTY*, *Sail_1164_G12*, was very helpful at establishing (or, in this case, confirming) the allelic relationships between all *RTY* gene mutants. This discovery emphasizes the critical importance of using true (i.e., full-length RNA) null alleles for all complementation tests, at least for the genes whose products may form dimers or higher order complexes and, therefore, interallelic complementation is possible.

Interallelic (also known as heteroallelic) complementation in transheterozygotes has been observed in many species, from yeast to humans, but is not a very commonly reported, nor widely known, phenomenon. In flowering plants, there is only a handful of cases attributed to interallelic complementation. Some of the first reports are in the *shrunken* sucrose synthetase (Chourey, 1971; Chourey and Nelson, 1979) and *aldehyde dehydrogenase* (Schwartz, 1975) loci of maize. The best-known examples of intragenic complementation in plants, however, come from the analysis of numerous *GNOM* alleles in Arabidopsis (Mayer et al., 1993; Busch et al., 1996; Grebe et al., 2000). *GNOM* encodes a guanine nucleotide exchange factor that regenerates ARF-GTPase involved in vesicle trafficking (Shevell et al., 1994, Busch et al., 1996). Mutations that abolish or severely reduce GNOM activity lead to dramatic defects in the establishment of apical-basal polarity during embryogenesis, at least in part due to the mislocalization of the auxin transporter PIN1 (Steinmann et al., 1999). Mayer and colleagues (Mayer et al., 1993) classified 24 different alleles of *GNOM* into three classes, A, B and C. A-and B-class mutants display at least partial inter-allelic complementation in transheterozygotes, whereas all other allelic combinations (i.e. within the A and B classes, between A and C alleles, or between B and C alleles) result in plants phenotypically similar to their parents. The ability of different A-and B-class *gnom* alleles to complement each other in transheterozygotes has been attributed to homomeric interactions between the GNOM subunits and/or partial stop-codon read-through in *gnom* nonsense alleles (Busch et al., 1996).

More recently, Goldraij and co-workers (Goldraij et al., 2003) reported interallelic complementation of the ubiquitous urease gene, known as *AJ6/EU4*, in soybean. De Leon and colleagues (de León <u>et al</u>., 2004) described a similar phenomenon for the cytokinin receptor *CRE1/WOL1* locus in Arabidopsis. Likewise, Shang and collaborators (Shang et al., 2011) observed interallelic complementation of the *BRI1* brassinosteroid receptor alleles in Arabidopsis that harbor mutations in extracellular and cytoplasmic (kinase-inactive) domains of the protein. Finally, Pysh (Pysh, 2015) reported intragenic complementation in transheterozygotes between two Arabidopsis alleles of the cellulose synthase gene, *AtCESA3*. Again, the common theme in the interpretation of all of these results is that the affected proteins function as dimers or higher order multimeric complexes: their ability to di- or multimerize leads to the complementation and, hence, restoration of missing activities in the complex. In fact, the Theologis group took advantage of this trans-subunit complementation phenomenon to explore functional homo- and heterodimerization between two tomato (Tarun et al., 1998) and nine Arabidopsis (Tsuchisaka et al., 2009) ACC synthase (ACS) family members by expressing pairs of recombinant ACS proteins with complementary mutations *in vitro* or in *E. coli* and accessing restoration of the ACS enzymatic activity.

Another example of interallelic complementation that does not require dimerization is found in genes coding for proteins that possess more than one enzymatic activity. For example, in *Nicotiana plumbaginifolia* mutations in the *nitrate reductase* gene that target different functional domains of the protein complement each other but not mutations that inactivate the same domain (Caboche and Rouzé, 1990). We have used this concept to functionally group the different alleles based on 3D structural information and the phenotypes of the transheterozygous mutant combinations. Theoretically, another way two mutant alleles of the same gene can generate a fully functional product in an interallelic combination is through mRNA trans-splicing (Mongelard et al., 2002; Lasda and Blumenthal, 2011). This mechanism splices together two or more mRNA molecules in trans, and therefore theoretically can “correct” mutant phenotypes of the missense alleles that are defective in different exons. Trans-splicing events for nuclear-encoded genes have been reported to occur in rice (Zhang et al., 2010; Mathew et al., 2016), maize (Lu et al., 2013) and tomato (Chen et al., 2019), but the frequency of these processing events appears to be much lower in plants than that in, for example, Drosophila (McManus et al., 2010). Thus, the physical separation of the different mutations in the cross-complementing *RTY* gene is consistent with three possibilities: complementation via protein dimerization, functional specialization of different protein domains, and mRNA trans-splicing. We do, however, favor a combination of the first two possibilities as these are supported by our computational modeling.

It is important to disclose here that multiple strong alleles of *RTY* have been previously isolated by different groups, yet no concerns about the interallelic complementation have been raised. The classical *rty* allele utilized as a reference in our study was first reported in a PhD thesis in 1994 (King, 1994) and published in late 1995 (King et al., 1995). In 1995, Celenza et al., reported the identification of a phenotypically similar *alf1-1* mutant (Celenza et al., 1995) and Boerjan et al., described seven additional alleles of this gene under the name of *superroot1 (sur1-1* through *sur1-7)* (Boerjan et al., 1995). In 1996, Lehman et al., published a *hls3* allele of *RTY* (Lehman et al., 1996). King and coauthors reported the results of complementation crosses between *rty* and *hls3*, as well as *rty* and two unpublished alleles of this gene, *invasive root-1 (ivr-1)* and *ivr-2*, with all three crosses revealing lack of complementation and, hence, allelism (King et al., 1995). Likewise, Celenza and colleagues stated lack of complementation in three crosses between *alf1-1, rty*, and *hls3* mutant, again suggesting that these mutations affect the same gene (Celenza et al., 1995). Finally, the seven *sur1* alleles have each been crossed to one other *sur1* allele and also found to be non-complementing (Boerjan et al., 1995). One obvious difference between these previously reported alleles of *RTY* and our new allelic series is the strength of the mutations. The EMS mutants described herein are phenotypically milder relative to the *rty* allele and are partially fertile (Fig. 3D), as these were derived from large M2 families from which all strong mutants (that were initially selected but failed to set seeds) have been simply lost. Unlike our missense alleles, many (but not all) of the previously published *RTY* alleles may be null. Nonetheless, given that the phenotypically null *rty* mutant could complement several of our weaker alleles of *RTY*, the cautionary take-home message of this study stays the same: allelism tests require the use of a full-length RNA-null mutant as a reference in complementation crosses or can otherwise provide misleading results. In our case, in light of the complementation observed in *rty* crosses, we chose to invest time into the characterization of these mutants because initially we erroneously concluded that these mutants were novel.

### Molecular modeling of dimeric RTY

The computational modeling of the three-dimensional *RTY* structure provided important clues to explain the F1 generation transheterozygote complementation in the RTY heterodimers and also demonstrated potential predictive power of the model. To our knowledge, this is the first time that these types of approaches have been used to structurally characterize transheterozygous mutations in plants. In other model organisms, the structural basis of interallelic complementation has been explored only in a handful of cases, such as for myosin light chain protein Cdc4p in *Schizosaccharomyces pombe* and acetylcholine receptor AChR in humans (Slupsky et al., 2001; Shen et al., 2016). We argue that combining genetics with structural modeling yields unique insights into the function of proteins harboring amino acid substitutions, as demonstrated by the characterization of mutant proteins encoded by the missense alleles of *RTY* and their interallelic combinations (see below).

Discerning structure-to-function relationships requires quality structural data (e.g., from nuclear magnetic resonance, X-ray crystallography, or cryo-electron microscopy) for the protein of interest or a related protein. To our knowledge, no protein structure is currently available for RTY from *Arabidopsis thaliana*, but several related proteins have been structurally characterized through crystallization. Although the RTY protein sequence aligns with tyrosine and other aromatic amino acid aminotransferases from mouse, human, and bacterial proteomes, we chose the sequence of MEE17 (PDB 5WMH) from *Arabidopsis thaliana*, a bifunctional aspartate aminotransferase and glutamate/aspartate-prephenate aminotransferase (Holland et al., 2018), as the template for our analysis. MEE17 is a far better template for this molecular modeling study, as it is derived from the same species as RTY, harbors PLP in the binding pocket, includes an extra alpha helical structure that is lacking in the related human tyrosine aminotransferase (PDB 3DYD) (Supplemental Fig. S4), contains coordinates for 399 out of 475 residues, and aligns well with the RTY sequence, with an E-value of 3.8E-17.

The structural interrogation of the RTY molecular model presented here allowed us to infer possible structure-to-function relationships for the mutations characterized in this study. We worked under the premise that mutations that affect the same functional domain should not complement each other in the transheterozygous plants. This basic assumption, together with the structural models, provides possible mechanism(s) for the disruption of the dimerization interface, cofactor binding pocket, or alteration of the catalytic domain. For example, D243 is likely directly involved in PLP/substrate binding and any disruption in this region is likely to interfere with RTY’s ability to recruit substrates. When examining the phenotypes of the transheterozygous F1 plants, we observed that G72D has the strongest phenotype when combined with the D243N mutation, suggesting that both affect the same functional domain, in this case, the PLP binding pocket. This is further supported by the fact that although G72 is not directly located in the cofactor binding pocket, it is a solvent-exposed residue that lines the channel entrance for this cofactor. Similarly, while the very different phenotypes of the P213 (strong) and G260 (weak) mutants do not suggest that these two mutations affect the same functional domain, their location immediately adjacent to the substrate-binding pocket and the fact that the strongest phenotype of any of the G260E-containing interallelic combinations is the one with P213S insinuates that these two mutations may disable the same protein functionality.

Residue D315 is positioned directly in the dimerization interface and any disruption here could lead to alteration of the interface and a decrease in the dimer’s ability to form. In addition, this mutation is also in close proximity to the substrate binding pocket of the adjacent monomer (∼8Å away). Of these two non-mutually exclusive possibilities, our analysis suggests that the D315N mutations predominantly affects dimerization and is less likely to affect substrate binding. This possibility is supported by the yeast two-hybrid data that revealed reduced binding of the mutant monomers to one another, as well as by the fact that D315N complemented all other missense mutations examined, including those predicted to affect the cofactor and substrate binding, thus disfavoring the possibility of D315N interfering with substrate recruitment. Conversely, the position of P404 in the 3D model failed to provide convincing support for a specific functional alteration in this mutant. Again, the combination of structural data analysis and transheterozygous phenotypes provided a reasonable hypothesis for the structure-function relationship of these mutations. In fact, distinct location of the P404S in the 3D model far away from the dimerization interface and the substrate binding pocket suggested that this mutation may disable a functional domain distinct from those affected by the other missense mutations analyzed. Consistent with this line of thought, we discovered partial complementation in all transheterozygotes containing the P404S mutations, suggesting that it impairs an activity not critically disturbed in any of the other alleles.

The 3D molecular model of RTY described above has provided structure-to-function reasons for the observed phenotypes for a majority of the *RTY* mutant alleles characterized in this study. We cannot rule out more nuanced structural reasons for the phenotypes observed, such as complementary effects of the mutations stabilizing either the substrate-binding pocket, dimerization interface, and/or interactions with potential partner proteins in some of the transheterozygous mutant combinations. Future experimental high-resolution structural data or molecular dynamic simulations could provide greater insight into the structural effects of these mutations, such as altered conformations within the dimerization interface leading to weaker dimers or within the substrate binding pocket reducing or abolishing substrate recruitment. In a meantime, a combination of genetics and 3D computational models enabled us to explain the molecular consequences of several *RTY* missense mutations and their interallelic combinations.

## Materials and Methods

### Mutant and complementation lines, growth conditions, and EMS mutagenesis

All of the new mutants described herein are in Columbia background. *rty* (CS8156)*, Sail_1164_G12* (CS878294), *sur2* (Salk_028573) and Col *DR5:GFP* (CS9361) were obtained from the Arabidopsis Biological Resource Center. Col *DR5-GUS* reporter (Ulmasov et al., 1997)*, YUC1* overexpression line (Zhao et al., 2001) and *cyp79b2 cyp79b3* T-DNA knockout (Zhao et al., 2002) were provided by Drs. Thomas Guilfoyle, Yunde Zhao, and John Celenza, respectively. *ctr1-1* (Kieber et al., 1993), *wei2-1, wei2-1 DR5:GUS, wei7-2, wei7-4, wei7-4 DR5:GUS, wei2-1 sur2, wei7-4 rty, wei2-1 rty* (Stepanova et al., 2005); *rty DR5:GUS* (Stepanova et al., 2007) and *sur2 cyp79b2 cyp79b3* (Stepanova et al., 2011) were previously reported. New mutant combinations were generated by crossing various mutants to each other and phenotyping/genotyping the lines in the F2 and F3 generation (see below). Reporter introgression was also done by crosses and phenotyping of progeny in F2 and F3. Seeds were germinated in plates in AT plates (1x Murashige and Skoog salts, 1% Sucrose, pH 6.0 with 1 M KOH, 0.7% Bacto-Agar) in the dark for 3 days and then moved to constant light, as indicated. For propagation, 10-to 14-day-old seedlings were transplanted to soil and grown under 16-hour light/8-hour dark cycle in 1:1 mix of Fafard superfine germinating mix and Fafard 4P mix.

A *RTY* cDNA complementation construct driven by the *35S* promoter and fused to a C-terminal TAP-tag in LIC6 vector was ordered from ABRC (DKLAT2g20610) and transformed into *TG8B* via floral dip method (Clough and Bent, 1998). T1s were selected in gentamycin and propagated, with images of complemented lines taken in the T2 generation.

EMS mutagenesis was performed on imbibed Columbia seeds as described (Guzmán and Ecker, 1990). The M1 generation was propagated in families of approximately 1000 plants each. A total of 27 M1 families were generated and screened in the M2 generation. M2 seeds were plated at 1000-2000 plants per 15cm Petri plate filled with 50ml of AT media (two plates per family), stratified in the cold for 2-4 days, germinated in the dark for 3 days and then transferred to light for additional 5-10 days. The plates were periodically screened visually for cotyledon epinasty. Putative mutants were picked, propagated in soil and re-tested in the M3 generation. One additional *RTY* allele, *IK6G12*, was derived from an equivalent EMS mutagenesis carried out in the *wei8-1* mutant background.

### PCR-based mutant mapping

Physical mapping of new mutants was performed in the F2 generation of crosses to L*er*. Epinastic F2 seedlings were picked from plates and transplanted to soil. A single leaf per plant was harvested, its genomic DNA extracted, and short genomic fragments that vary in length between Col and L*er* were PCR-amplified in 10uL reactions (1uL genomic DNA, 1 uL 10xPCR buffer, 0.25uL 2 mM dNTPs, 0.25uL homemade Taq polymerase, 0.25 mL 20uM forward primer, 0.25uL 20uM reverse primer, and 7uL H2O) using 40 cycles of the following PCR program: 30 s at 94°C, 30 s at 56°C, 1 min at 72°C. PCR products were separated on 3-4% agarose 1xTAE gels, photographed, and scored. The indel PCR marker primers F26H11-1-F1 and F26H11-1-R1 utilized in mapping of *RTY* mutants are listed in Table S1.

### Mutant genotyping

Plant genotyping was performed by CEL1 assay and/or Sanger sequencing (for the EMS mutants) and by PCR (for T-DNA mutants). PCR primers are listed in Supplemental Table S1. PCR conditions were the same as described for mapping, except longer amplification times were allowed (1min per Kb).

For the CEL1 assay, gene-specific primers were used to amplify overlapping gene fragments from the mutant and wild-type Columbia samples in 20ul PCR reactions (2uL genomic DNA, 2 uL 10xPCR buffer, 0.5uL 2 mM dNTPs, 0.5uL homemade Taq polymerase, 0.5 mL 20uM forward primer, 0.5uL 20uM reverse primer, and 14uL H2O) using 40 cycles of the following PCR program: 30 s at 94°C, 30 s at 56°C, 1-2 min [1min per kb] at 72°C. 5ul of each mutant and 5ul of the wild-type sample for each fragment were combined, and then denatured and annealed to each other in a Thermocycler by heating the mixture for 10 min at 99°C, reducing the temperature to 70°C, and running 70 steps of the program reducing the temperature 0.3°C per step, keeping the samples for 20 seconds at each temperature, going from 70°C to 49°C. Annealed 10ul samples were combined with 10ul CEL1 mix (2ul 10xCEL1 buffer, 1ul CEL1, 7ul diH2O), incubated at 45°C for 15-20 minutes, and then immediately moved to ice. Digested chilled samples were resolved on a 1% agarose 1xTAE gel by gel-electrophoresis, photographed and scored.

### GUS staining and basic GFP microscopy

Samples harboring the *DR5:GUS* transgene were harvested in ice-cold 90% acetone and stored at −20C overnight or longer. To stain for GUS, samples were washed once with Rinse Buffer (50mM Sodium Phosphate buffer, pH7, 0.5mM K3Fe(CN)6, 0.5mM K4Fe(CN)6) and stained overnight in Rinse Buffer supplemented with 1ug/ml X-gluc (with X-gluc dissolved in DMSO at 5mg per 100ul and diluted with Rinse Buffer to 1ug/ml final concentration). Staining reactions were stopped with 15% ethanol. To remove chlorophyll, GUS-stained seedlings were incubated for 2-3 days in 70% EtOH at 37-50°C. Samples were photographed using Q Capture software with a Q Imaging digital camera hooked to a Leica dissection scope. To visualize expression of *DR5:GFP*, a Zeiss Axioplan epifluorescence microscope was utilized. Images were captured with a Diagnostic Instruments Color Mosaic camera using Spot Insight software.

### Yeast two-hybrid assay

To test the level of interaction between RTY monomers, we employed the yeast two-hybrid assay. Binding between wild-type monomers, TG8B mutant monomers, and wild type with TG8B monomers was evaluated. The cDNA clone G10872 (ABRC) was employed as RTY-WT and used as template for the PCR-based approach to generate the TG8B mutant ORF. To obtain TG8B, three PCR reactions (iProof, BioRad) were performed: PCR 1 using primers SUR1_For and sur1_RevTG8B_Internal; PCR 2 with sur1_ForTG8B_Internal and SUR1_Rev (No STOP); and the fusion PCR 3 to join together both fragments harboring the mutation was conducted using the external primers SUR1_For and SUR1_Rev (No STOP) (see Supplemental Table S1 for primer information). The final PCR product was subcloned into pENTR/D-Topo. Both WT and TG8B RTY were transferred to pACT-GW (GAL4-Activation Domain, AD) and pAS-GW (GAL4-Binding Domain, BD) (Nakagawa et al., 2007; Shimoda et al., 2008) by Gateway LR clonase recombination. The Y190 yeast strain was transformed with the resulting “bait” constructs, harboring the GAL4-BD. Strains were tested for lack of trans-activation and then, transformed with the “prey” constructs containing the GAL4-AD. Interaction between “bait” and “prey” was inferred from the activity of the *LacZ* reporter (Deplancke et al., 2006). Qualitative colony-lift assays were performed to assess the activity of β-galactosidase by visual scoring of the levels of the resulting blue staining in the presence of X-gal (Goldbio) substrate (Deplancke et al., 2006). To quantify the β-galactosidase activity, lysed yeast cells were resuspended in Z-buffer (Deplancke et al., 2006), supplemented with O-nitrophenyl-beta-D-galactopyranoside, ONPG (Sigma), and the amount of yellow product obtained from the ONPG catalysis was measured by determining the absorbance at 420 nm (Smale, 2010) using a SpectraMax M2 microplate reader.

### Analysis of conserved residues in the RTY protein

Conservation of the amino acid residues mutated in our allelic series of RTY was analyzed using the Multiple Sequence Alignment (MSA) tool from Phytozome (Goodstein et al., 2012; https://phytozome.jgi.doe.gov) by comparing RTY-like proteins of the *Brassicales* and *Malvales* orders of flowering plants from the following species: *Arabidopsis halleri*, *Arabidopsis lyrata*, *Arabidopsis thaliana*, *Boechera stricta*, *Brassica oleracea*, *Brassica rapa*, *Capsella grandiflora*, *Capsella rubella*, *Carica papaya*, *Eutrema salsugineum*, *Gossypium raimondii*, and *Theobroma cacao*. Utilizing the MSA results, a sequence logo was generated to represent the level of conservation of each mutated residue by employing WebLogo (Crooks et al., 2004; http://weblogo.berkeley.edu/logo.cgi).

### RNA extraction and qRT-PCR

8-day-old seedlings of the indicated genotypes grown for 3 days in the dark followed by 5 days in constant light were collected into 1.5 ml Sarstedt screw-cap tubes prefilled with four 2.4mm Zirconia/Silica beads each, frozen in liquid nitrogen, and ground in a Vivadent shaker. For *rty* and *Sail_1164_*G12 mutants that are sterile, homozygous seedlings were phenotypically selected from the progeny of heterozygous parents. Total RNAs were extracted (Rueber and Ausubel, 1996) and reverse transcribed with random hexamer primers using a TaqMan reverse transcription kit (Applied Biosystems). SYBR Green qPCRs were performed in an Applied Biosystems StepOnePlus Real-Time PCR System (Thermo Fisher Scientific) with Power SYBR green Master Mix (Applied Biosystems) using primers For1RTY-qRT and Rev1RTY-qRT for *RTY* and At5g44200F and At5g44200R for the control gene, *At5g44200* (see Supplemental Table S1 for primer information). Normalized mean values for *RTY* levels plus/minus standard deviation were plotted on a bar graph.

### Molecular modeling

MODELLER v9.18 was used to construct a model of the dimeric *RTY* wild-type and transheterozygotes using *Arabidopsis thaliana* prephenate aminotransferase (PDB: 5WMH; Holland et al., 2018). During the model building process, we employed an optimization method involving conjugate gradients and molecular dynamics to minimize violations of the spatial restraints. 500 models were generated from an alignment of the Arabidopsis sequence and scored by the internal MODELLER scoring method Discrete Optimized Protein Energy (DOPE; Shen and Sali, 2006). The structure with the lowest DOPE score was subsequently run through PROCHECK and WHATCHECK (checks the stereochemical quality of a protein structure) for quality (Laskowski et al., 1993; Hooft et al., 1996). All images were produced with PyMOL (Baker et al., 2001; The PyMOL Molecular Graphics System, Version 2.0 Schrödinger, LLC.).

## Acknowledgements

We would like to thank Jeonga Yun for technical assistance with sequencing *RTY* alleles and C. Douglas Grubb for helpful discussions on interallelic complementation. This work was supported by the NSF grants MCB0923727, IOS1444561 and IOS1650139 to A.N.S. and J.M.A, MCB1158181 to J.M.A, and IOS1650139 to A.N.S.

---

**Supplemental Figure S1.**
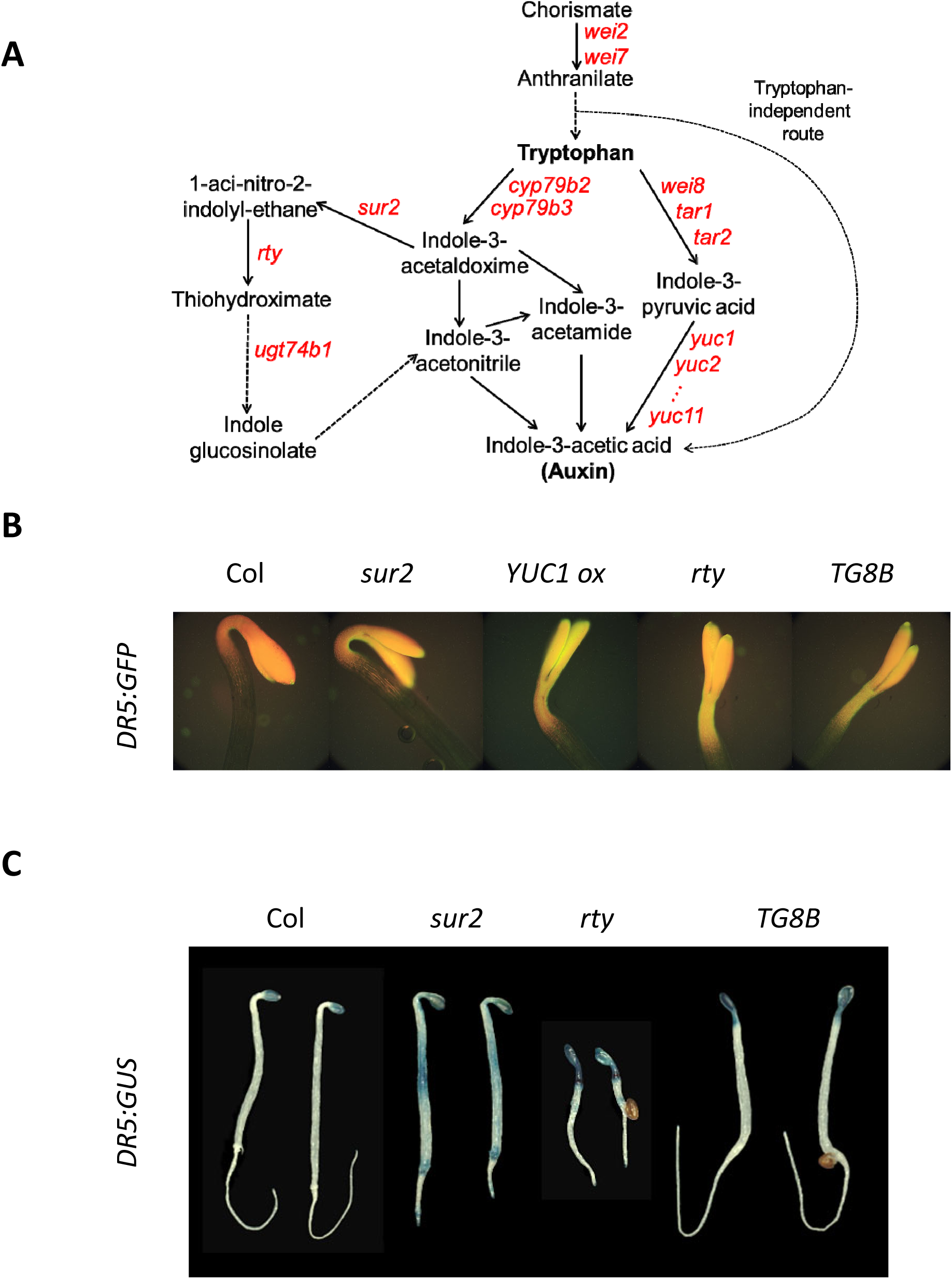
Increased levels of auxin responses in auxin overproducing mutants. (A) A model of the auxin biosynthesis pathway displaying positions of auxin mutants utilized or referenced in this study. Metabolic intermediates are marked in black and mutants in the genes that encode the enzymes catalyzing specific steps are shown in red. Solid arrows represent single reactions, whereas dashed arrows correspond to multiple enzymatic steps. (B, C) The activity of *DR5:GFP* (B) and *DR5:GUS* (C) reporters is elevated in high-auxin mutants.

**Supplemental Figure S2.**
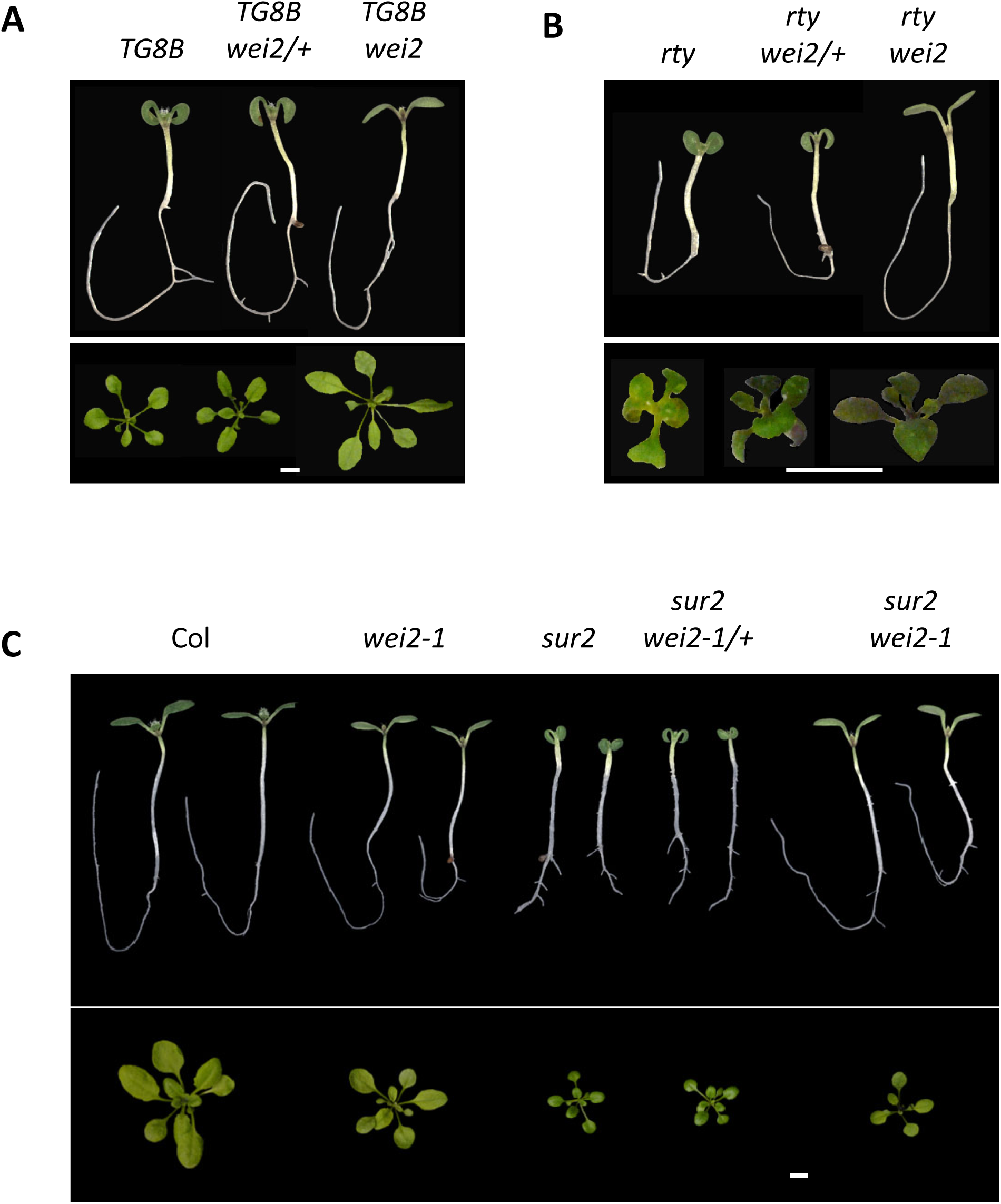
Lack of suppression of the *TG8B, rty*, and *sur2* defects by one copy of *wei2-1*. 8-day-old seedlings grown in plates for 3 days in the dark followed by 5 days in the light are shown in upper panels (A, B, C). 4-week-old soil-grown (A, C) and 3-week-old plate-grown (B) plants are displayed in lower panels. Scale bars in the lower panels represent 5 mm.

**Supplemental Figure S3.**
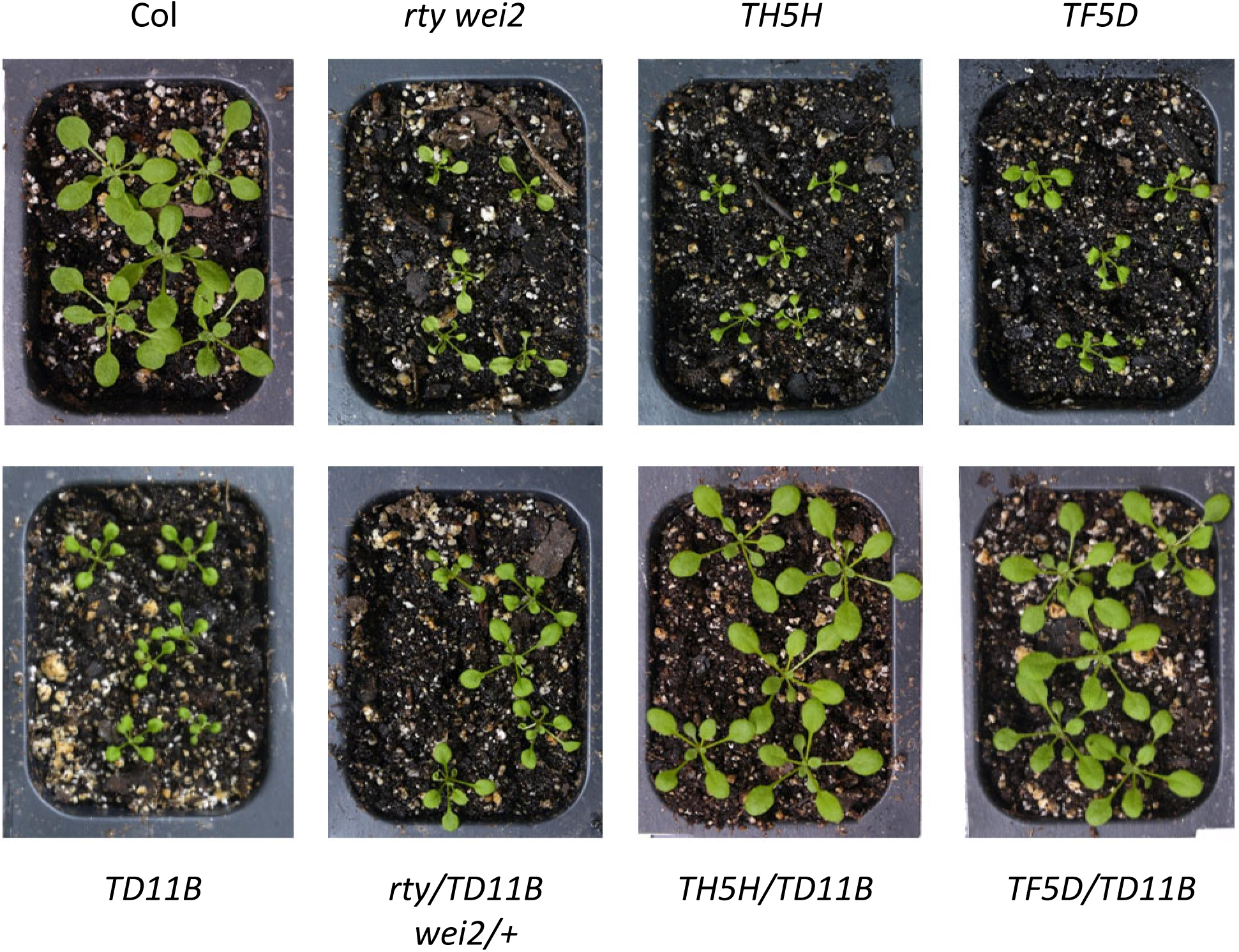
Interallelic combinations of *RTY* mutants. Examples of different degrees of partial complementation in 4-week-old soil-grown F1 plants are displayed: marginal complementation in *rty/TD11B wei2/+* and near complete recovery in *TH5H/TD11B* and *TF5D/TD11B* are observed.

**Supplemental Figure S4.**
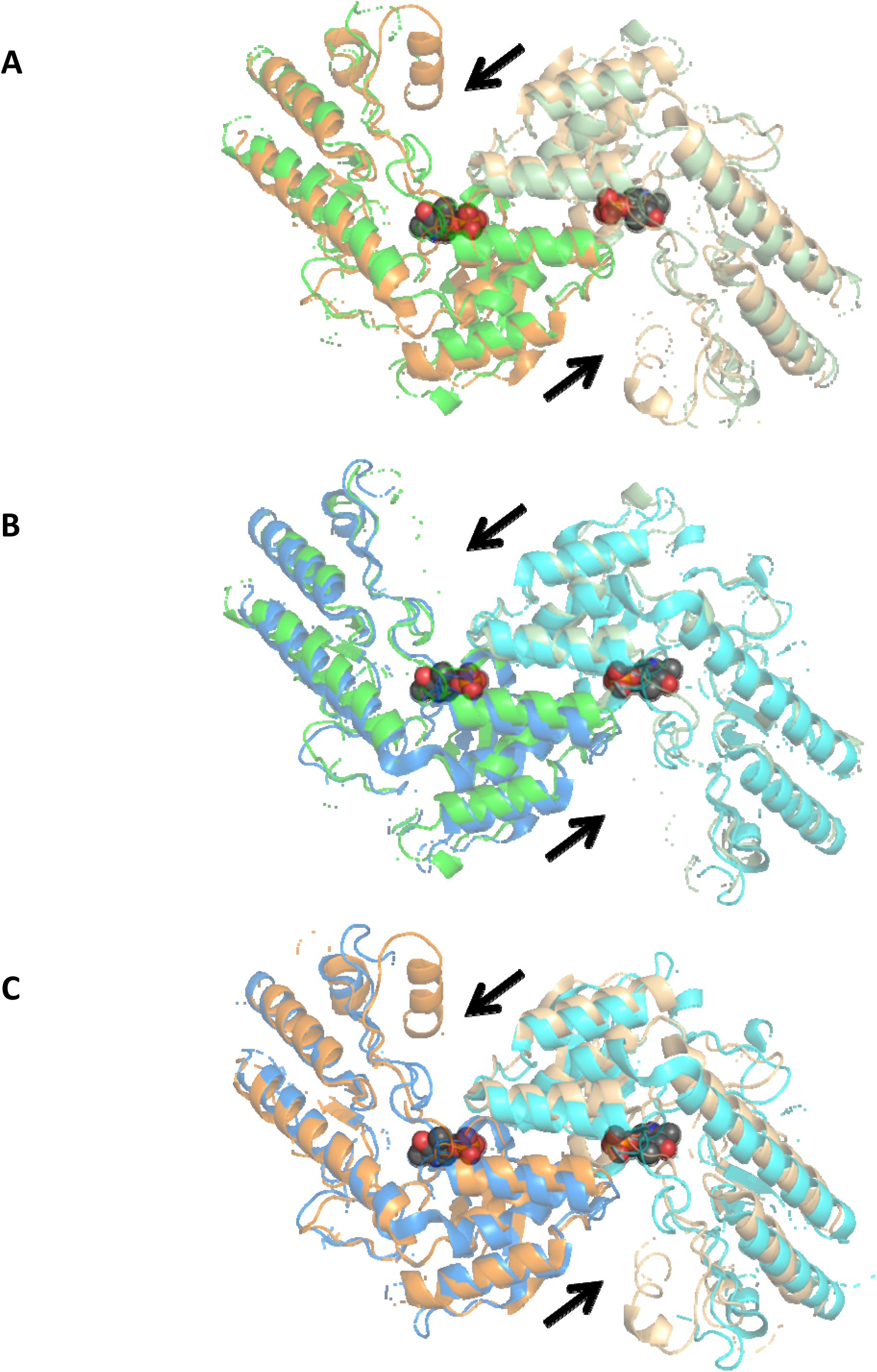
The crystal structure of the *Arabidopsis thaliana* bifunctional aspartate aminotransferase and glutamate/aspartate-prephenate aminotransferase MEE17 (PDB 5WMH) was used as a template to produce the RTY model. (A) Overlay of the generated RTY model (green and pale green cartoons) with MEE17 (orange and beige cartoons). (B) Overlay of the RTY model (green and pale green cartoons) with the human tyrosine aminotransferase TAT (PDB 3DYD, blue and cyan cartoons). (C) Overlay of MEE17 (orange and beige cartoons) with TAT (blue and cyan cartoons). PLP substrate is shown in sphere format. An extra alpha helical structure present in Arabidopsis RTY and MEE17 monomers but lacking in human TAT is labeled with black arrows for clarity.

**Supplemental Figure S5.**
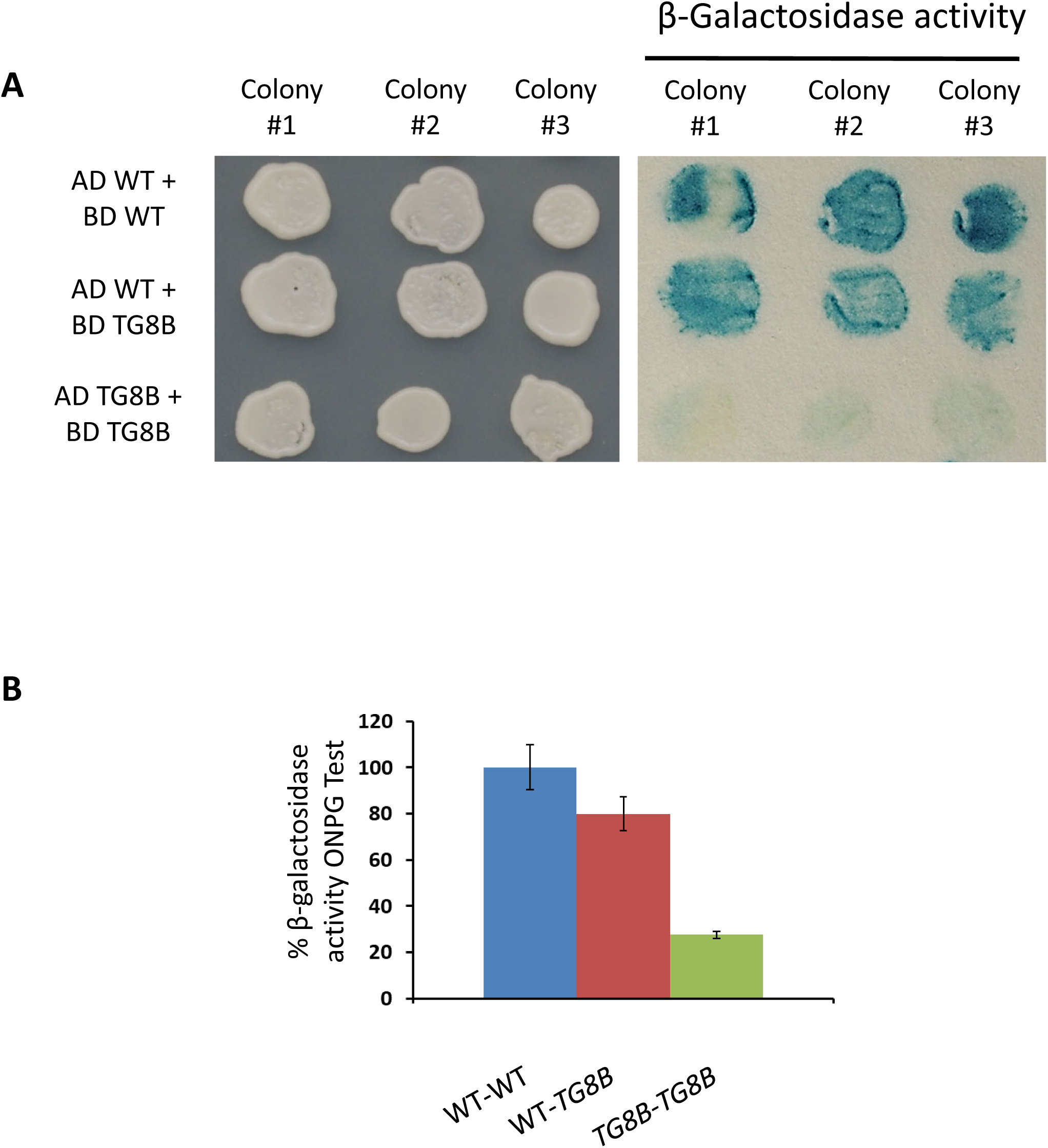
Reduced dimerization capacity of the *TG8B* (D315N) version of RTY in yeast. Yeast two-hybrid assay was employed to assess the ability of RTY proteins to dimerize. (A) Qualitative β-galactosidase activity assay in plates. (B) Quantification of β-galactosidase activity in liquid assay employing ONPG as substrate. Each bar is an average of two different experiments, including three biological replicates and five technical replicates per experiment. Mean relative β-galactosidase activity values plus/minus standard deviation are displayed.

**Supplemental Figure S6.**
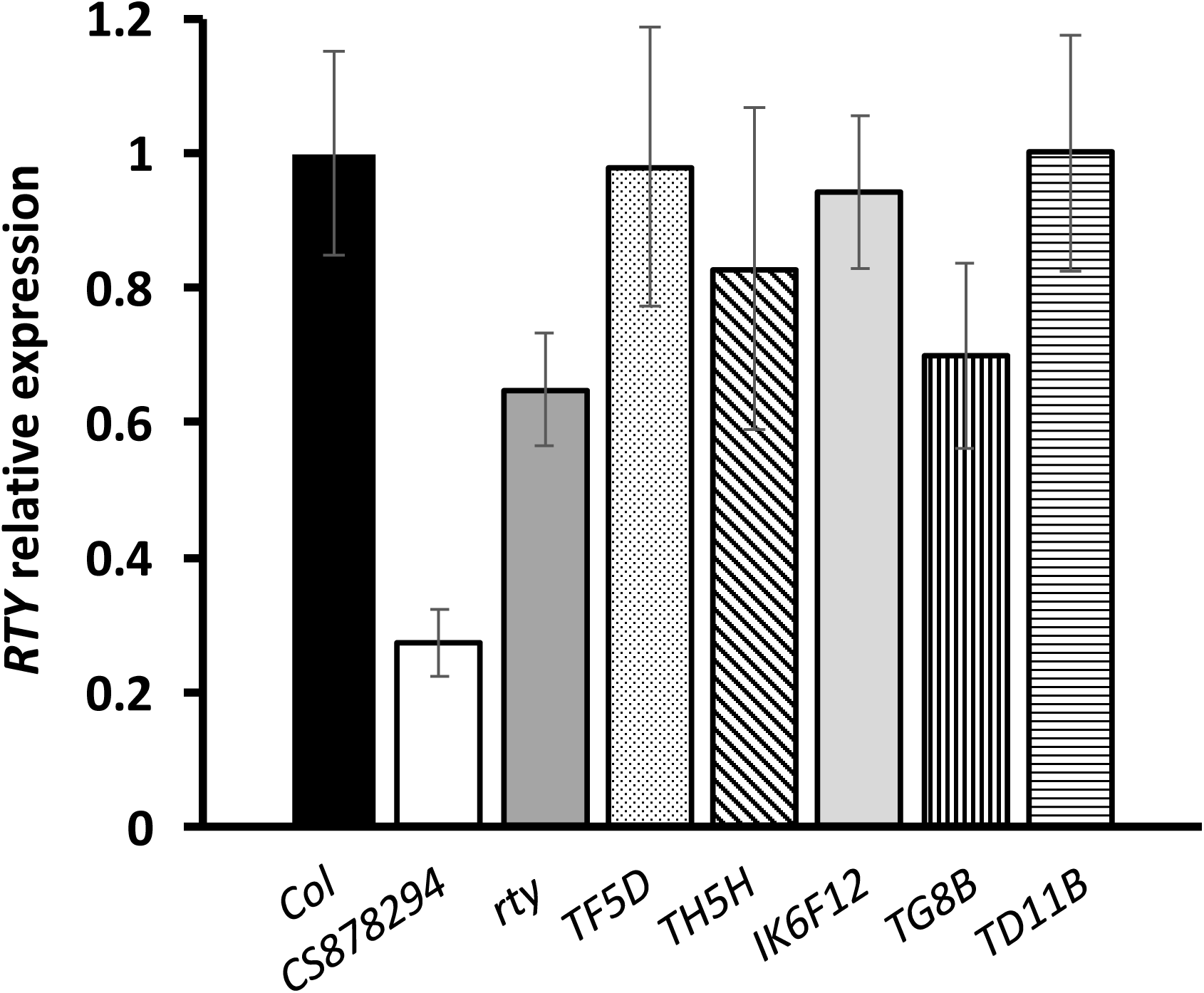
Quantification of *RTY* expression in various alleles of *RTY* by qRT-PCR. Three biological and three technical replicates were tested per genotype. Relative expression of *RTY (At2g20610)* was plotted relative to the control gene, *AtCBP80 (At5g44200).* The bars correspond to the mean relative expression values plus/minus standard error of the mean.

**Supplemental Table Sl.**
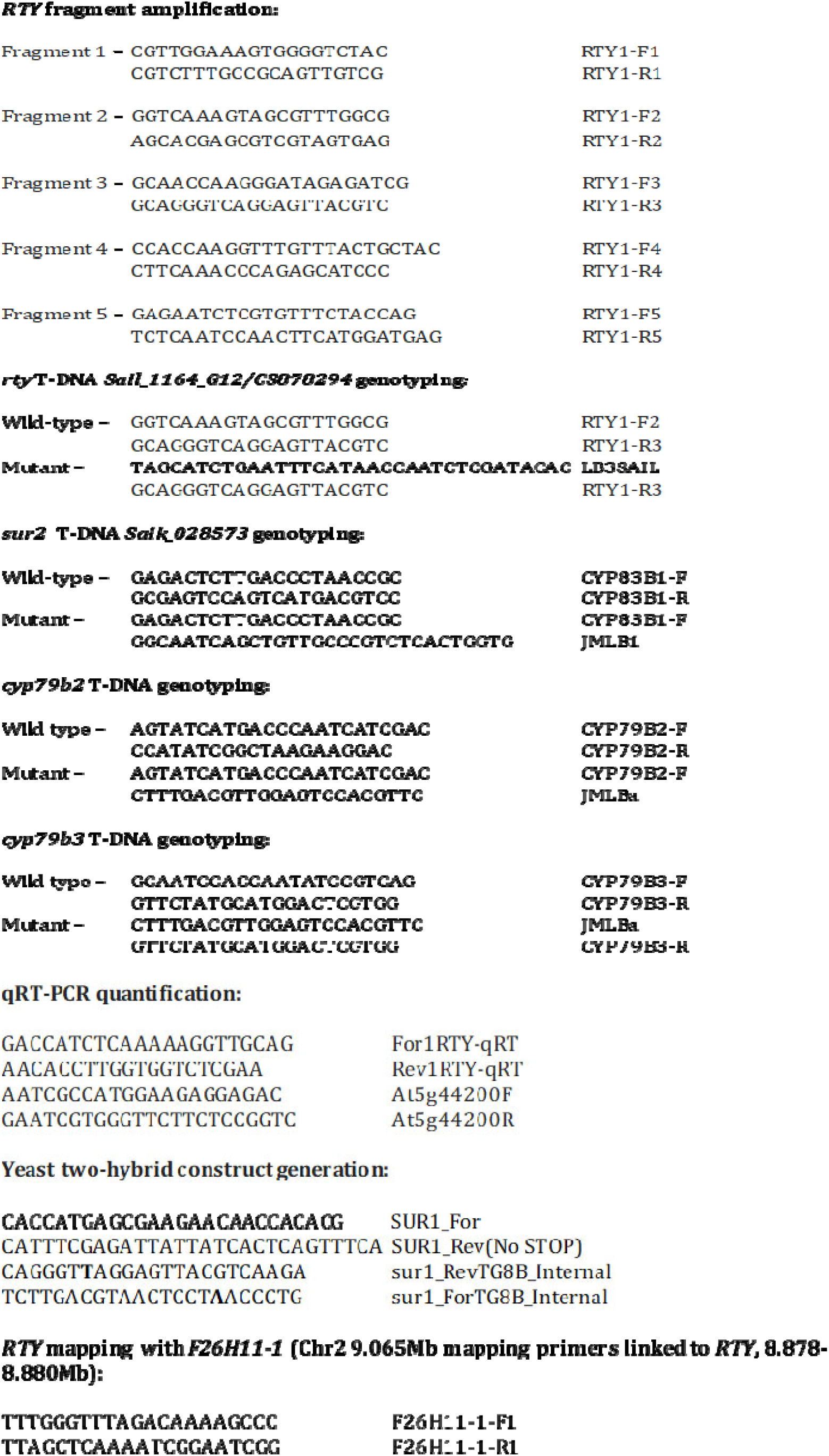
Primers employed in this study

